# A rapid phylogeny-based method for accurate community profiling of large-scale metabarcoding datasets

**DOI:** 10.1101/2022.12.06.519402

**Authors:** Lenore Pipes, Rasmus Nielsen

**Affiliations:** Department of Integrative Biology, University of California-Berkeley, Berkeley, California, USA; GLOBE Institute, University of Copenhagen, Copenhagen, Denmark

**Author notes:** Correspondence and requests for materials should be addressed to Rasmus Nielsen.

## Abstract

Environmental DNA (eDNA) is becoming an increasingly important tool in diverse scientific fields from ecological biomonitoring to wastewater surveillance of viruses. The fundamental challenge in eDNA analyses has been the bioinformatical assignment of reads to taxonomic groups. It has long been known that full probabilistic methods for phylogenetic assignment are preferable, but unfortunately, such methods are computationally intensive and are typically inapplicable to modern Next-Generation Sequencing data. We here present a fast approximate likelihood method for phylogenetic assignment of DNA sequences. Applying the new method to several mock communities and simulated datasets, we show that it identifies more reads at both high and low taxonomic levels more accurately than other leading methods. The advantage of the method is particularly apparent in the presence of polymorphisms and/or sequencing errors and when the true species is not represented in the reference database.

---

In the past ten years, metabarcoding and metagenomics based on DNA sequencing and subsequent taxonomic assignment, have become an important approach for understanding diversity and community organization at many taxonomic levels. This has led to the publication of over 80 taxonomic classification methods^1^. There are three major strategies in classification methods: (1) composition-based, which do not align sequences but extract compositional features (e.g., kmers) to build models of probabilistic taxonomic inclusion, (2) alignment-based, which rely on alignments to directly compare query sequences to reference sequences but do not use trees, and (3) phylogenetic-based, which rely on a phylogenetic tree reconstruction method, in addition to alignments, to perform a placement of the query onto the tree. As a trade-off between speed and precision for processing Next-Generation Sequencing (NGS) data, the vast majority of recent classification methods have either relied on alignment-based or composition-based strategies.

Composition-based tools reduce the reference database by indexing compositional features such as kmers for a rapid search of the database. These methods require an exact match between the kmer in the query sequence and the kmer in the reference database. As a result of hash indexing of kmers, kraken2^2^, for example, can classify *>*1 million reads within 1 minute using the entire Geengenes or SILVA databases^3^. Alignment-based tools use a fast local aligner such as BLAST^4^ to pairwise align queries to the reference database, and define a score based on sequence similarity in the alignment between the read and reference sequence. However, alignment-based methods can be many orders of magnitude slower than composition-based tools since datasets with *>*10 million reads require weeks of BLASTN running time^5^. In both composition-based tools and alignment-based tools, a lowest common ancestor (LCA) algorithm is then typically used to assign at different taxonomic levels (**Figure 1A**). LCA works by assigning to the smallest possible clade that include all matches with a similarity less than the specified cut-off.

**Figure 1:**
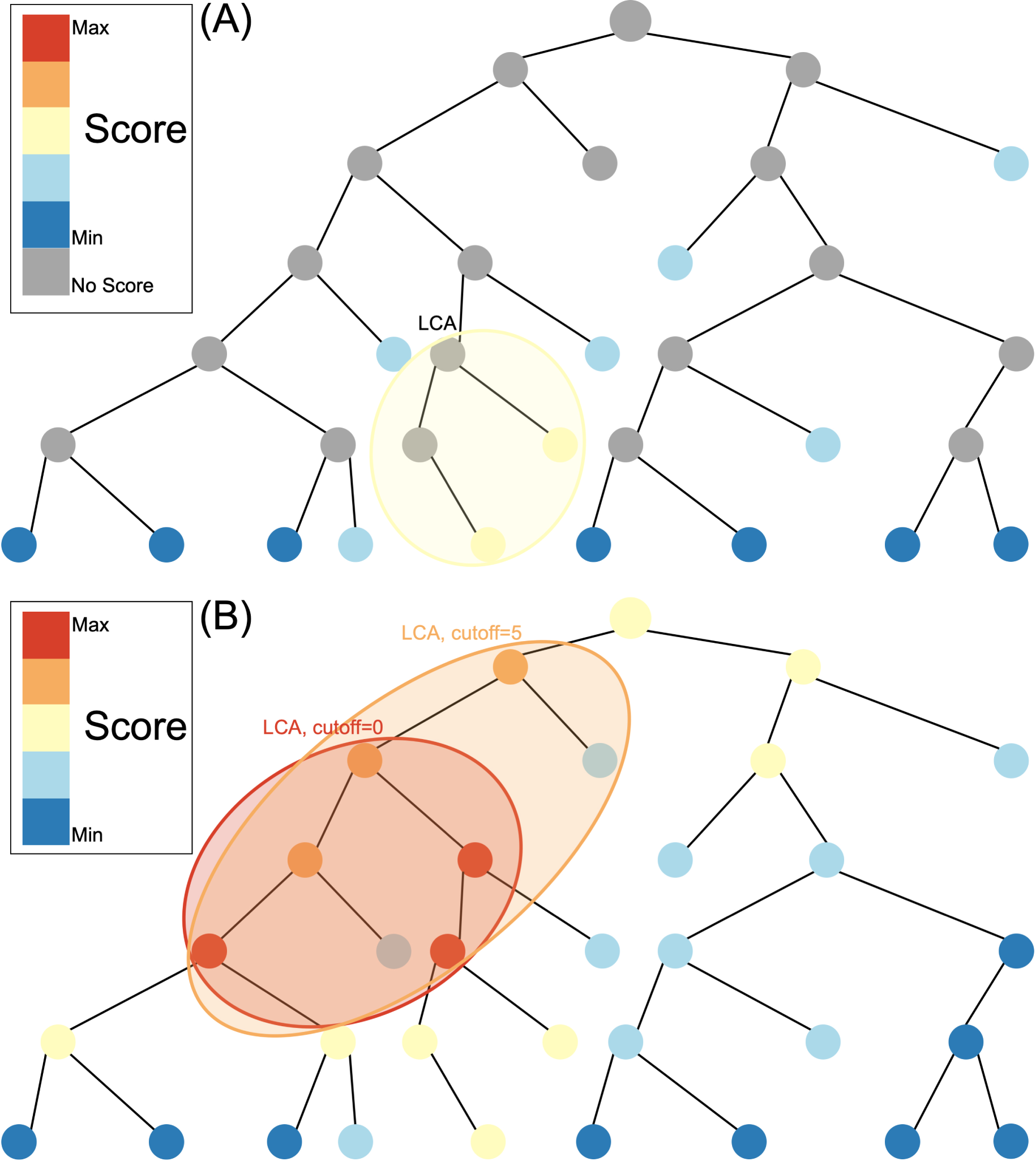
Species assignment in alignment-based methods (A) vs. Tronko (B). In Tronko, scores are calculated for all nodes in the tree based on the query’s global alignment to the best `BWA-MEM` hit. The query is assigned to the LCA of the highest scoring nodes within the cut-off threshold.

Phylogenetic placement methods place a query sequence onto a phylogenetic tree of reference sequences. This placement requires a full multiple sequence alignment (MSA) of the reference sequences and a subsequent estimation of a phylogenetic tree. However, large datasets with high rates of evolution are hard to align accurately^6^ and phylogeny estimation methods produce poor trees when MSAs are not of high quality^7^. Futhermore, phylogentic placement tends to be computationally demanding as both running time and memory usage scale linearly with the size of the reference database ^8^. Even for reference databases that contain sequences as few as 1,600 sequences, assignment for a single query using the most cited phylogenetic placement method, *pplacer*^9^, takes more than 7 minutes and requires over 10GB of RAM (on a Dell PowerEdge server with 32 CPU threads and 512GB of RAM). At this rate, a reference database that contains a metabarcode such as Cytochrome oxidase 1 (COI) that has at least 1.5 million reference sequences, assigning just a single query would require 20.9 hours and 2.37TB RAM. Scaling the query size to millions of queries would therefore be computationally intractable.

To address these challenges, the most recent implementations of phylogeny-based methods^10^ rely on reference database reduction techniques (i.e., using only representative taxa or consensus sequences for a sparse backbone tree) to handle the large amount of data that is routinely produced. Often a single species is selected to represent an entire clade^11^. While this reduces the computational cost, it also reduces the granularity, and potentially the accuracy, of the assignments. As a trade-off between speed and precision, the vast majority of recent classification methods are either alignment-based or composition-based approaches^12^ since phylogeny-based methods have not scaled to handle the entirety of the rapidly growing reference databases of genome markers and the increasingly large amounts of NGS data.

Here we describe a new method for phylogenetic placement, implemented in the program ‘Tronko’ (https://github.com/lpipes/tronko, **Supplementary Software**), the first phylogeny-based taxonomic classification method designed to truly enable the use of modern-day reference databases and NGS data. The method is based on approximating the phylogenetic likelihood calculation by (1) only allowing the edge connecting the reference sequence to the tree to join at existing nodes in the tree and then (2) approximating the likelihood using a probabilistically weighted mismatch score based on pre-calculated fractional likelihoods stored in each node (See Methods). We argue that (2) approximates the full maximized likelihood assignment without requiring any numerical maximization under the approximating assumption that the read joins the tree in an existing node with a zero length branch. The approximation is equivalent to calculating the expected average mismatch to each node in the phylogeny. The assignment method in Tronko uses the LCA criteria but, unlike composition-based and alignment-based approaches (**Figure 1A**), takes advantage of fractional likelihoods stored in all nodes of the tree with a cut-off that can be adjusted from conservative to aggressive (**Figure 1B**). In the simplest case, when the reference sequences form a single tree, Tronko uses a pre-calculated MSA, the phylogenetic tree based on the MSA, and precalculated posterior probabilities, which are proportional to the fractional likelihoods. However, in more typical cases, when a single tree/MSA is unsuitable for analyses, as the reference sequences encompass increasingly divergent species as well as an increasing volume of sequences, we present a fully customizable divide-and-conquer method for reference database construction that is based on dividing reference sequences into phylogenetic subsets that are re-aligned and with local trees re-estimated.

The construction of the database, MSAs, and trees, facilitates fast phylogenetic assignment. The assignment algorithm then proceeds by (1) A BWA-MEM ^13^ search on all sequences in the database, (2) a pairwise sequence alignment between the query and the top hit in each alignment-subset containing a BWA-MEM hit using either the Needleman-Wunsch algorithm^14^ or the Wave-front Alignment algorithm^15^, and (3) a calculation of a score based on the approximate likelihood for each node in subsets with a BWA-MEM hit. An additional LCA assignment for all subsets can then be applied to summarize the results. For full details, please see METHODS.

## RESULTS

To compare the new method (Tronko) to previous methods, we constructed reference databases for COI and 16S for common amplicon primer sets using CRUX^16^ (See Methods for exact primers used). We first compared Tronko to pplacer and APPLES-2 for reference databases containing a reduced amount of sequences (*<*1,600 sequences) to compare the speed and memory requirements with comparable phylogenetic-based assignment methods. Tronko shows speed-ups *>*20 times over pplacer, with a vastly reduced memory requirement illustrating the computational advantage of the approximations in Tronko (**Supplementary Fig. S8**). Tronko demonstrates a speed-up *>*2 times over APPLES-2 with a similar memory footprint. In terms of accuracy all methods had a 100% true positive rate at the species level. Additionally, in terms of the species assignment rate (the percentage of queries that were assigned at the species level), Tronko assigns the most queries.

Next, in addition to pplacer and APPLES-2, we evaluated Tronko’s performance to kmer-based kraken2^2^ which previously has been argued to have the lowest false-positive rate^3^, and two other popular alignment-based methods: MEGAN^17^ and metaphlan2^18^. We used two types of cross validation tests: leave-one-species-out and leave-one-individual-out analyses. The leave-one-species-out test involves removing an entire species from the reference database, simulating next generation sequencing reads from that species, and then attempting to assign those reads with that species missing from the database. The leave-one-individual-out test involves removing a single individual from the reference database, simulating next generation sequencing reads from that individual, and then attempting to assign those reads with that individual missing from the database. In both tests, singletons (i.e., cases in which only one species was present in a genera or cases in which only one individual represented a species) were exempt from the tests.

We performed a leave-one-species-out test comparing Tronko (with LCA cut-offs for the score of 0, 5, 10, 15, and 20 with both Needleman-Wunsch alignment and Wavefront alignment) to kraken2, metaphlan2, and MEGAN for 1,467 COI sequences from 253 species from the order Charadriiformes using 37,515 (150bp*×*2) paired-end sequences and 768,807 single-end sequences (150bp and 300bp in length) using 0, 1, and 2% error/polymorphism (**Figure 2**). We use the term “error/polymorphism” to represent a simulated change in nucleotide that can be either an error in sequencing or a polymorphism. We display confusion matrices to display the clades in which each method has an incorrect assignment (**Figure 3**). See **Figure S2** for results with the Wavefront alignment algorithm^15^.

**Figure 2:**
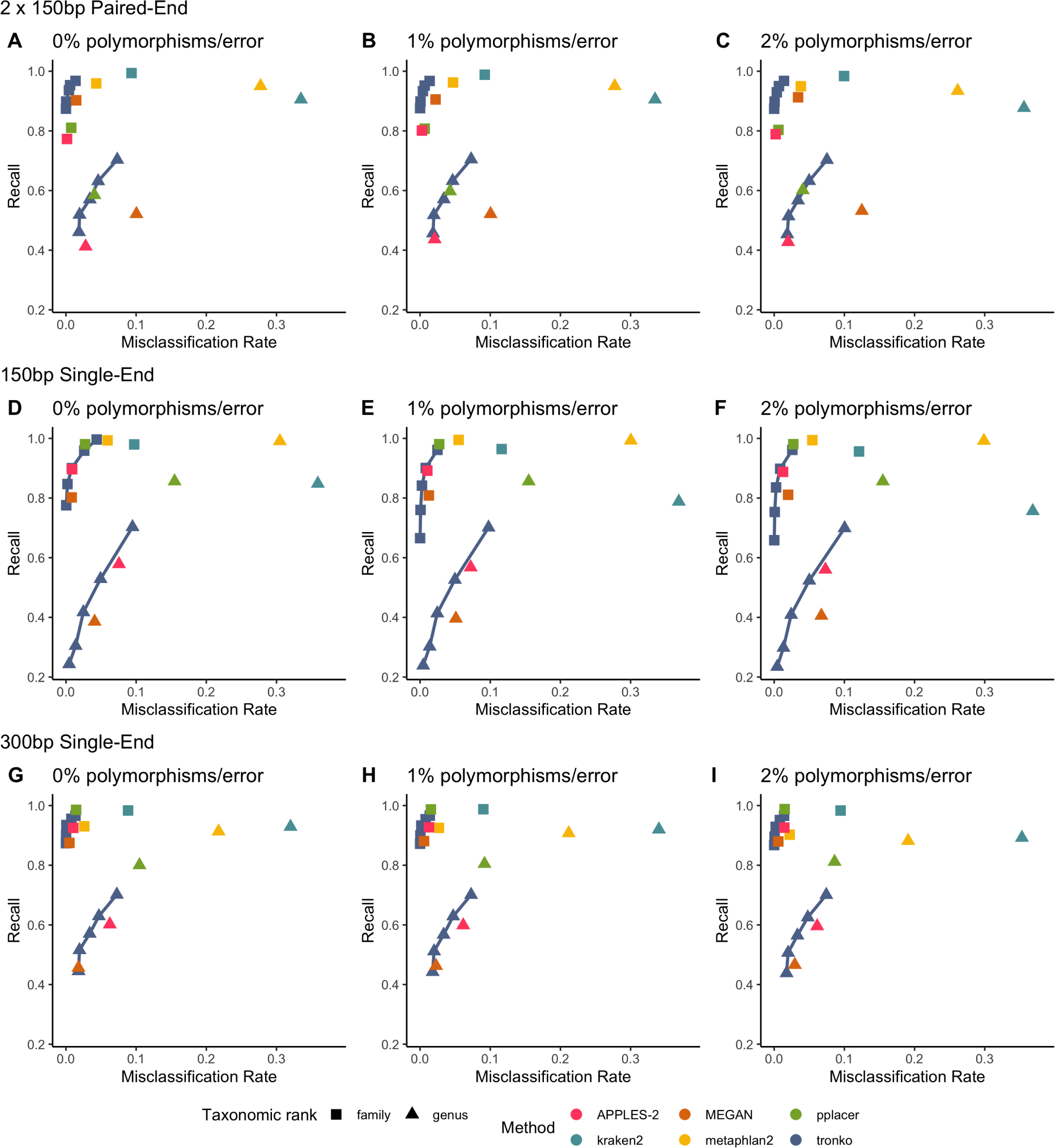
Recall vs. Misclassification rates using leave-one-species-out analysis of the order Charadriiformes (COI metabarcode) with paired-end 150bp*×*2 reads with 0% (A), 1% (B), and 2% (C) error/polymorphism, single-end 150bp reads with 0% (D), 1% (E), and 2% (F) error/polymorphism, and single-end 300bp reads with 0% (G), 1% (H), and 2% (I) error/polymorphism using kraken2, metaphlan2, MEGAN, pplacer, and APPLES-2, and Tronko with cut-offs of 0, 5, 10, 15, and 20 using the Needleman-Wunsch alignment (solid line).

**Figure 3:**
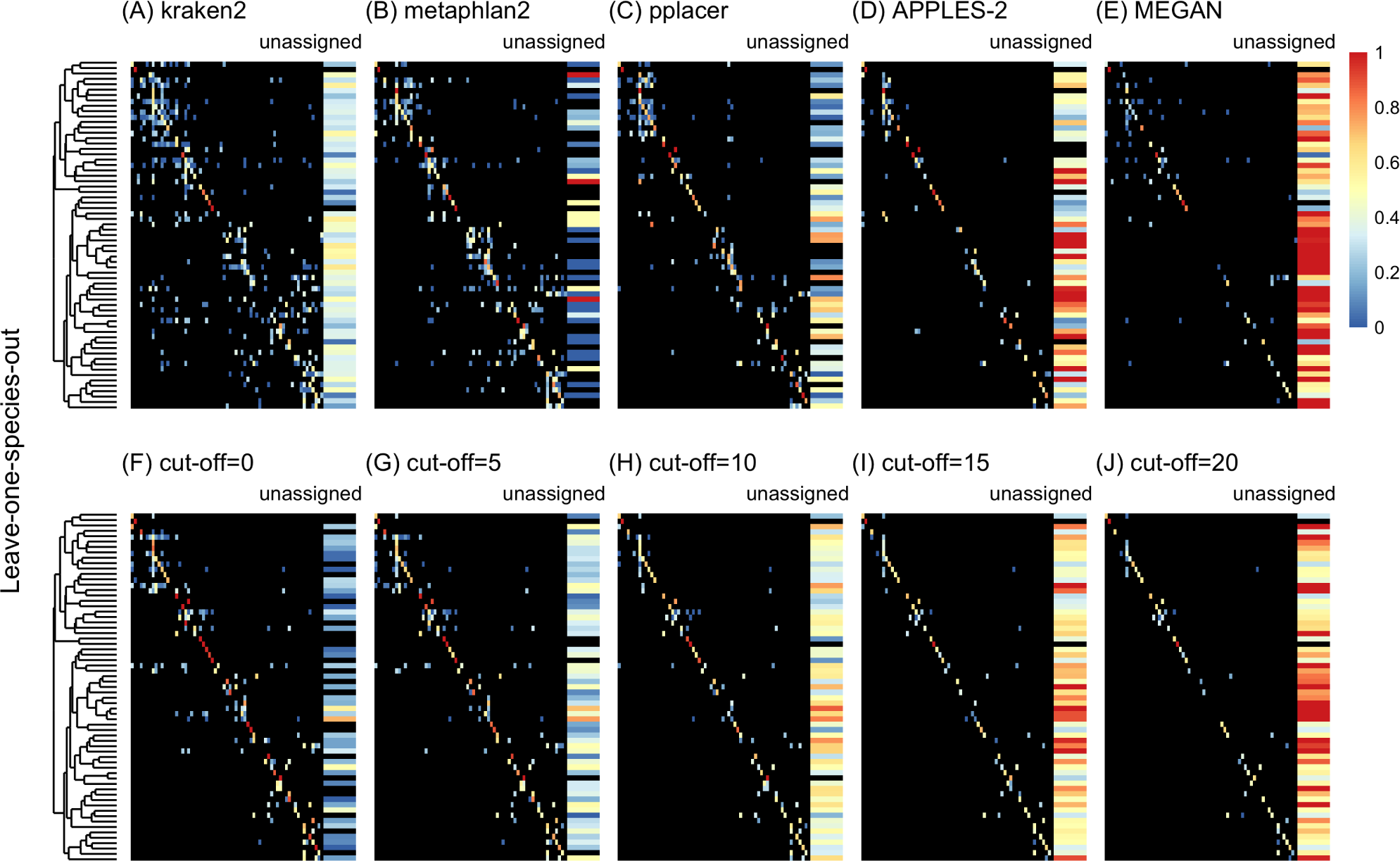
Confusion matrices at the genus level of the order Charadriiformes (COI metabar-code) using the leave-one-species-out analysis with paired-end 150bp*×*2 reads with 2% error/polymorphism using kraken2 (A), metaphlan2 (B), pplacer (C), APPLES-2 (D), MEGAN (E), and Tronko using the Needleman-Wunsch alignment (NW) for cut-offs 0 (F), 5 (G), 10 (H), 15 (I), and (J) 20. Unassigned column contains both unassigned queries and queries assigned to a lower taxonomic level. Phylogenetic tree represents ancestral sequences at the genus level.

Using leave-one-species-out and simulating reads (both paired-end and single-end) with a 0-2% error (or polymorphism), Tronko detected the correct genus more accurately than the other methods even when using an aggressive cut-off (i.e., when cut-off=0) (**Figure 3F**). Using 150bp paired-end reads with 1% error, Tronko had a misclassification rate of only 9.8% with a recall rate of 70.1% at the genus level using a cut-off set to 15 while kraken2, MEGAN, and metaphlan2 had misclassification rates of 33.5%, 10.0%, and 27.7%, respectively, with recall rates of 90.6%, 52.1%, and 95.0% (see **Figure 2B**). Tronko had a lower misclassification rate relative to the recall rate out of all methods for 150bp *×* 2 paired-end reads with 0% error/polymorphism (**Figure 2A**), 1% error/polymorphism (**Figure 2B**), and 2% error/polymorphism (**Figure 2** and **Figure 3D-I**), for 150bp reads with 0% error/polymorphism (**Figure 2D**), 1% error/polymorphism (**Figure 2E**), and 2% error/polymorphism (**Figure 2F**), and for 300bp reads with 0% error/polymorphism (**Figure 2G**), 1% error/polymorphism (**Figure 2H**), and 2% error/polymorphism (**Figure 2I**). See Methods for definitions of recall and misclassification rates. Tronko also accurately assigned genera from the Scolopacidae family (top left of matrices in **Figure 3**) using Needleman-Wunsch with a cut-off of 10 compared to kraken2, metaphlan2, and pplacer.

Next, we performed a leave-one-individual-out test for the same COI reference sequences using 746,352 single-end reads and 36,390 paired-end reads (**Figures 4** and **S4F-J**). See **Figure S5** for results with Wavefront alignment algorithm^15^. Using single-end reads of lengths 150bp and 300bp, Tronko has a lower misclassification rate and higher recall rate than kraken2, metaphlan2, and MEGAN. Using 150bp paired-end reads with 0% error (**Figure 4D**), Tronko had a misclassification rate at only 0.1% with a recall rate of 58.6% at the species level using a cut-off set to 10 while kraken2, MEGAN, and metaphlan2 had misclassification rates of 1.5%, 0.1%, and 11.0%, respectively, with recall rates of 85.4%, 60.7%, and 98.14%. Both metaphlan and kraken2 have a number of mis-assignments within the family of Laridae (see blue points across the diagonal in **Figure S4A** and **B**) and Tronko is able to accurately assign species within this family or assign at the genus or family level. We also observe that for increasing error rates, kraken2 and metaphlan2 have a substantial increase in misclassification rate. We believe it would be slightly misleading to display results for pplacer and APPLES-2 here due to the lack of an implementation to calculate the LCA on similar likelihoods. See **Figure S5** for results for pplacer and APPLES-2 along with Wavefront alignment algorithm.

**Figure 4:**
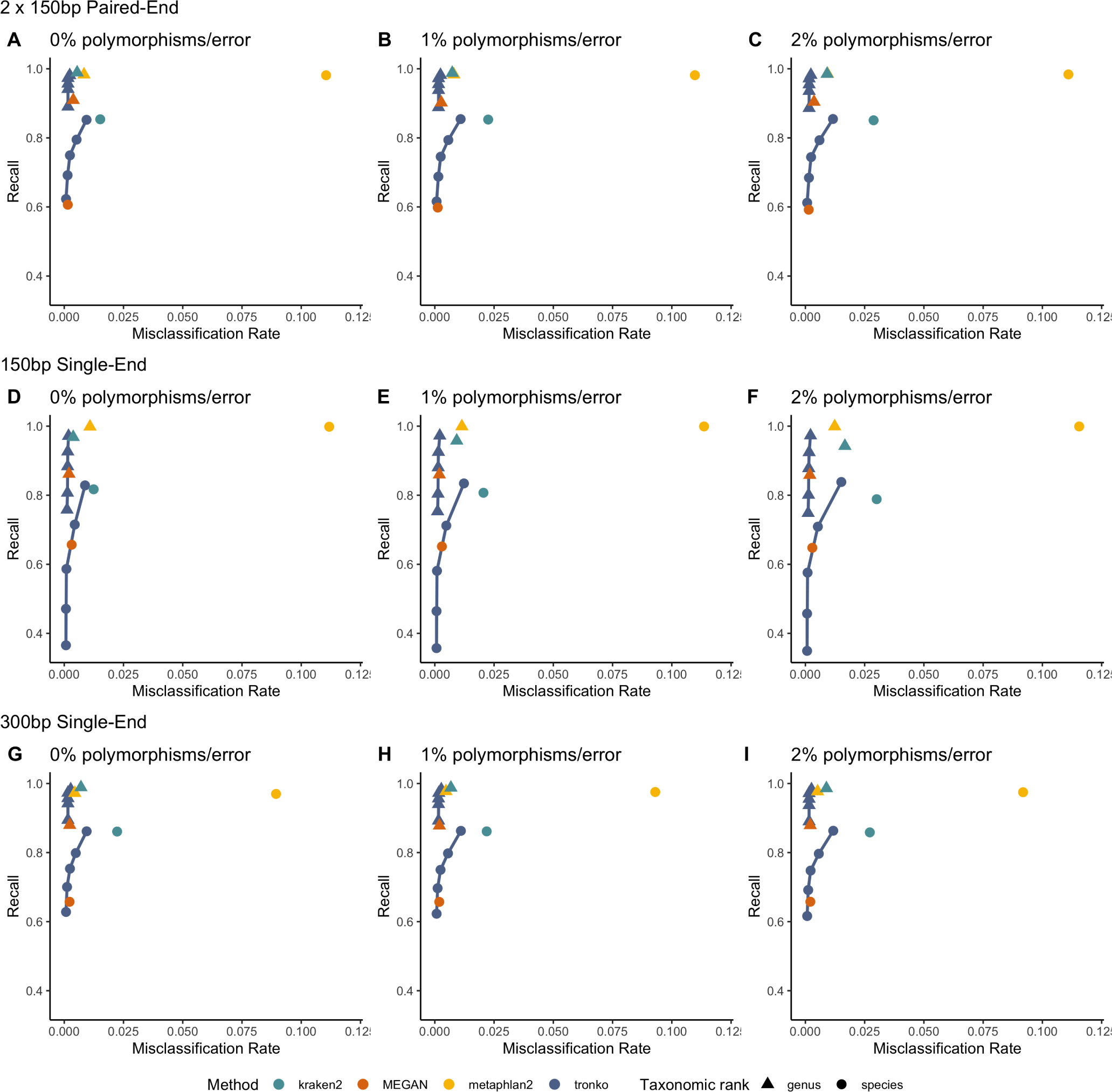
Recall vs. Misclassification rates using leave-one-individual-out analysis for the order Charadriiformes (COI metabarcode) with paired-end 2*×*150bp reads with 0% (A), 1% (B), and 2% (C) error/polymorphism, single-end 150bp reads with 0% (D), 1% (E), and 2% (F) error/polymorphism, and single-end 300bp reads with 0% (G), 1% (H), and 2% (I) error/polymorphism using kraken2, metaphlan2, MEGAN, pplacer, APPLES-2, and Tronko with cut-offs of 0, 5, 10, 15, and 20 using the Needleman-Wunsch alignment (solid line).

In order to replicate real-world scenarios, we added a leave-one-species-out test using 16S from 2,323 bacterial species and 5,000 individual sequences (**Figure 5**). We selected the sequences for the 16S dataset by grouping the sequences by the class level in a random order, rotating that order, and randomly selecting an individual sequence from each group. We then simulated sequencing reads from the dataset simulating 21,947,613 single-end reads (150bp and 300bp in length) as well as 21,478,738 paired-end 150bp*×*2 reads. In all simulations at both the genus and family levels, Tronko had a higher recall and a lower misclassification rate than all other methods. The simulations for 300bp single-end reads are not directly comparable to the 150bp single-end or paired-end reads because only 105 missing-out tests out of 2,310 were able to be performed because most reference sequence were *<*300bp in length. We only display the results for 300bp single-end reads for APPLES-2 in the supplement, as we believe the results are not a good representation of the method. See **Figure S6** for results for APPLES-2 using 300bp single-end reads along with results using the Wavefront alignment algorithm. Additionally, we tested the use of hmmer or MAFFT for alignments with APPLES-2 and pplacer (**Figure S7**), and we did not observe any substantial difference with the choice of alignment.

**Figure 5:**
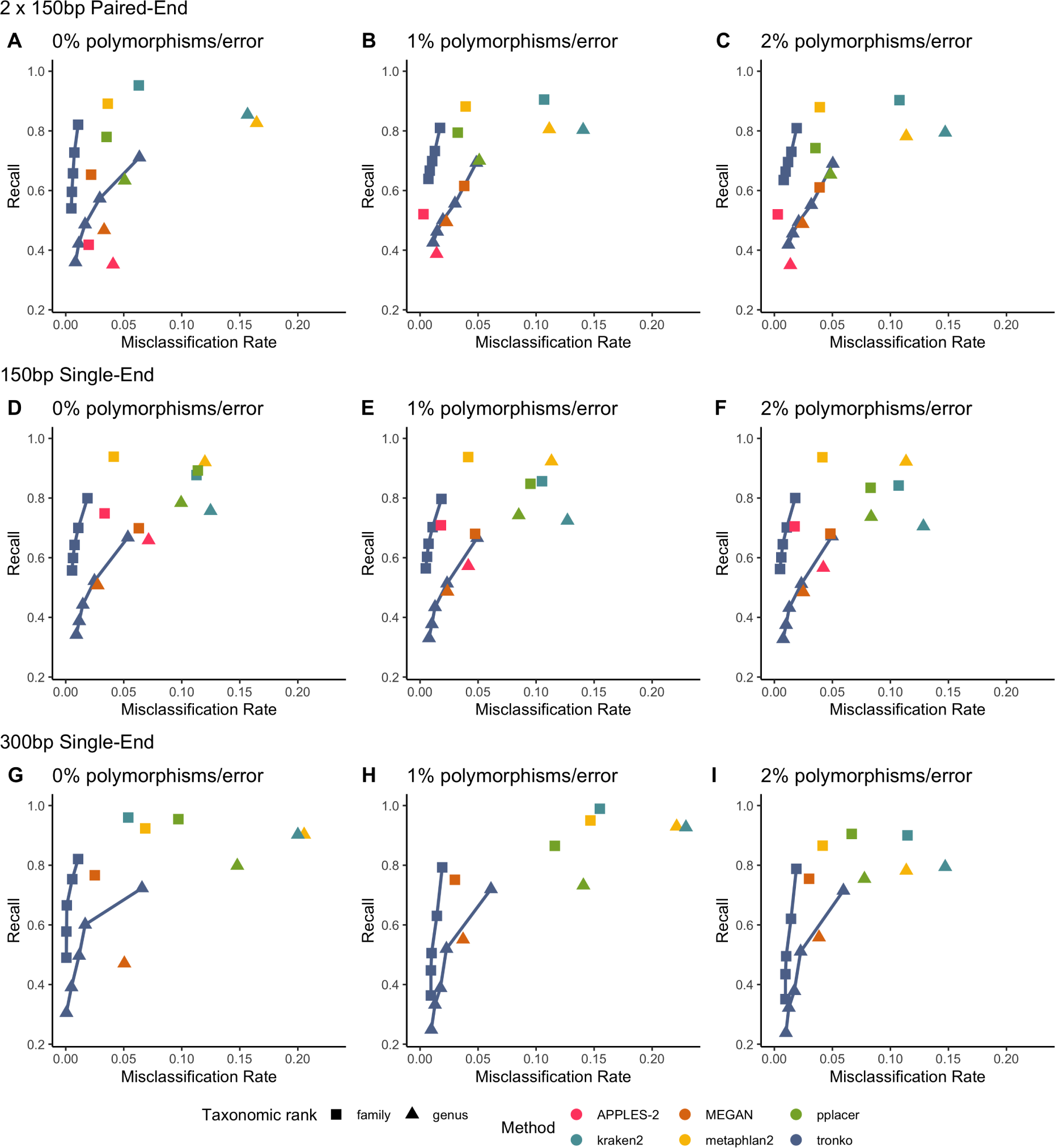
Recall vs. Misclassification rates using leave-one-species-out analysis with bacteria species (16S metabarcode) with paired-end 150bp*×*2 reads with 0% (A), 1% (B), and 2% (C) error/polymorphism, single-end 150bp reads with 0% (D), 1% (E), and 2% (F) error/polymorphism, and single-end 300bp reads with 0% (G), 1% (H), and 2% (I) error/polymorphism using kraken2, metaphlan2, MEGAN, pplacer, APPLES-2 and Tronko with cut-offs of 0, 5, 10, 15, and 20 using the Needleman-Wunsch alignment (solid line).

We then compared Tronko’s performance to kraken2, MEGAN, and metaphlan2 using mock communities for both 16S^19, 20^ and COI markers^21^ (**Figure 6**). We did not compare mock community data to pplacer and APPLES-2 because we were unsuccessful in building a full multiple sequence alignment for our 16S and COI reference databases. Tronko also relies on sequence alignments, but as described in the methods section, they can be handled by dividing sequences into clusters in the Tronko pipeline. For 16S, we used three different mock community datasets. We used 1,054,868 2*×*300bp Illumina MiSeq sequencing data from a mock community consisting of 49 bacteria and 10 archaea species from Schirmer *et al*. (2015)^19^, 54,930 2*×*300bp Illumina MiSeq sequencing data from a mock community consisting of 14 bacteria species from Lluch *et al*. (2015)^22^, and 206,696 2*×*300bp Illumina MiSeq sequencing data from a mock community of 20 evenly distributed bacterial species from Gohl *et al*. (2016)^20^. For the data from Schirmer *et al.* (2015), at the species level, Tronko had a less than 0.6% misclassification rate at every cut-off with a recall rate of 11.0% at cut-off 0 (**Figure 6A**; See **Figure S9** for plot without outliers). kraken2 had a misclassification rate of 1.2% with a recall rate of 10.6% when using its default database, and a misclassification rate of 3.5% and a recall rate of 35.1% when using the same reference sequences as Tronko. metaphlan2 did not have any assignments at the species, genus, or family level using the default database, and it had an 8.3% misclassification and 8.9% recall rate at the species level when using the same reference sequences as Tronko. MEGAN had a recall rate of 0.2% and a misclassification rate of 0% at the species level.

**Figure 6:**
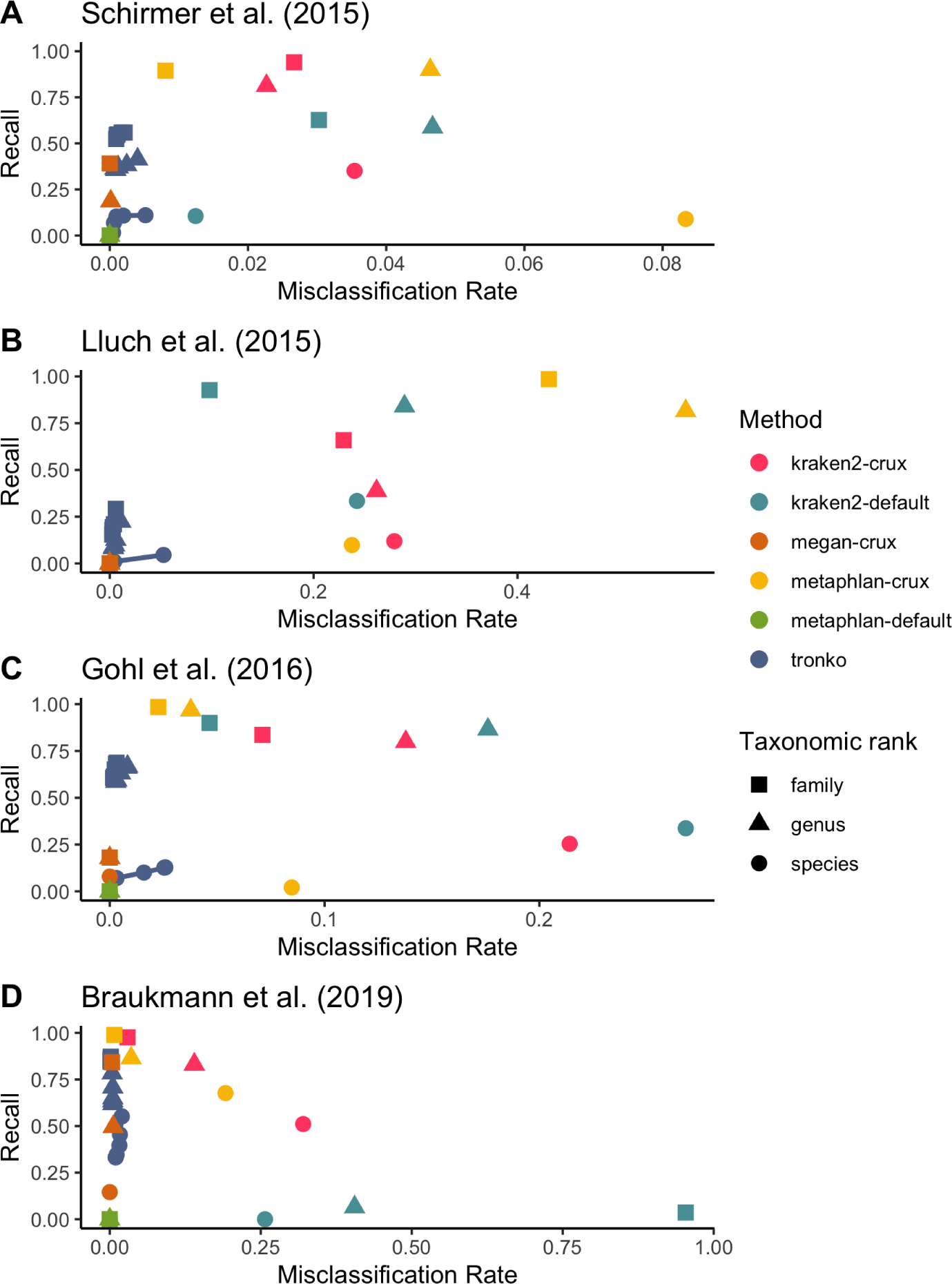
Recall vs. Misclassification rates using mock communities from Schirmer et al. (2015)^19^(A), Lluch et al. (2015)^22^(B), Gohl et al. (2016)^20^(C), and Braukmann et al. (2019)^21^(D) using both Needleman-Wunsch and Wavefront alignment algorithms. Figures with smaller misclassification rates on the x-axis are available for Schirmer et al. (2015), Lluch et al. (2015), Gohl et al. (2016), Braukmann et al. (2019) in Supplementary Figures S9, S10, and S12, respectively.

For the data from Lluch *et al.* (2015), at the genus level, Tronko had a misclassification rate of 0.6% and a recall rate of 22.3% using a cut-off of 0, while all other methods had a misclassification rate of *>*8% (see **Figure S10** for a close-up of the rates).

For the data from Gohl *et al.* (2016), at the species level, Tronko had a less than 2.6% misclassification rate at every cut-off with a recall rate of 12.8% at cut-off 0 (**Figure 6C**; See **Figure S11** for plot without outliers). kraken2 had a misclassification rate of 26.8% and recall rate of 33.7% when using its default database, and a misclassification rate of 21.4% and recall rate of 25.4% when using the same reference sequences as Tronko. metaphlan2 did not have any assignments at the species, genus, or family level using the default database, and it had an 8.5% misclassification and 2.1% recall rate at the species level when using the same reference sequences as Tronko. MEGAN had a misclassification rate of 0% and a recall rate of 4.4% at the species level.

For COI, we used a dataset from Braukmann et al. (2019)^21^ which consists of 646,997 2*×*300bp Illumina MiSeq sequencing data from 374 species of terrestrial arthropods, which is the most expansive mock community dataset that we used. At the genus level, Tronko had a misclassification rate of less than 0.5% with a recall rate of 78.3% at the cut-off of 0 (**Figure 6D**; see **Figure S12** for plot without outliers). With the default database, kraken2 had a misclassification rate of 40.5% with a recall rate of 6.5%. With the same reference sequences as Tronko, kraken2 still had a misclassification of 14.0% with a recall rate of 83.1%. metaphlan2 had a misclassification rate of 3.5% with a recall of 86.4% with the same reference sequences as Tronko while the default database failed to assign any reads. MEGAN had a 15.0% recall and 0% misclassification rate at the species level and a 49.9% recall and 0.5% misclassfication rate at the genus level.

We compared Tronko with kraken2, metaphlan2, and MEGAN (using BLAST as the aligner) for running time (**Figure 7**A) and peak memory (**Figure 7B**) using 100, 1,000, 10,000, 100,000, and 1,000,000 sequences using the COI reference database. Unsurprisingly, kraken2 had the fastest running time followed by metaphlan2, but MEGAN had a substantially slower running time than all methods. Tronko was able to assign 1,000,000 queries in *∼*8 hours with the choice of aligner being negligible. Tronko had the highest peak memory (*∼*50GBs) as it stores all reference sequences, their trees, and their posterior probabilities in memory. We note that for very large databases, the memory requirements can, in theory, be reduced by processing different alignment subsets sequentially.

**Figure 7:**
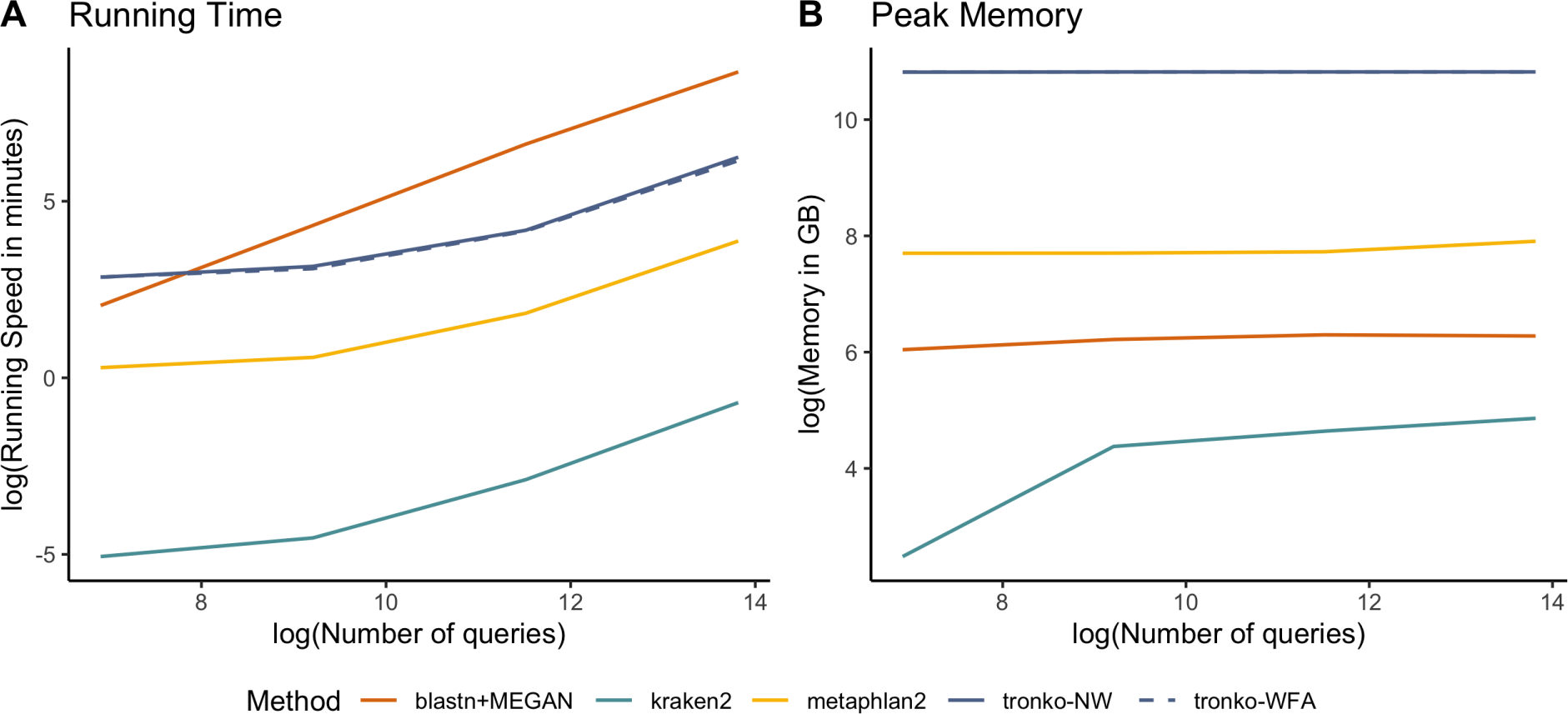
Comparisons of running time (A) and peak memory (B) using 100, 1000, 10000, 100000, and 1000000 queries for Tronko, blastn+MEGAN, kraken2, and metaphlan2 using the COI reference database. Needleman-Wunsch is NW and Wavefront alignment is WFA.

## Discussion

Both leave-one-species out and leave-one-individual-out simulations show that Tronko recovers the correct taxonomy with higher probability than competing methods and represents a substantial improvement over current assignment methods. The advantage of Tronko comes from the use of limited full sequence alignments and the use of phylogenetic assignment based on a fast approximation to the likelihood.

We evaluate Tronko using diferent cut-offs representing different trade-offs between recall and misclassification rate, thereby providing some guidance to users for choice of cut-off. We note that in most cases, the other methods evaluated here fall within the convex hull of Tronko, showing that Tronko dominates those methods, and in no cases do other methods fall above the convex hull of Tronko. However, in some cases other methods are so conservative, or anti-conservative, that a direct comparison is difficult. For example, when using single-end 300bp reads (**Figure 4G-I**), MEGAN has assignment rates that are so low that a direct comparison is difficult.

Among the methods compared here, kraken2 is clearly the fastest (**Figure 7A**). However, it generally also has the worst performance with a higher misclassification rate than other methods, especially in the leave-one-species out simulations (**Figure 2**).

Both metaphlan2 and MEGAN tend fall within the convex hull of Tronko. Typically, metaphlan2 assigns much more aggressively, and therefore, has both a recall and misclassification rate that is much higher than MEGAN, which assigns very conservatively. We also note that the computational speed of MEGAN is so low that it, in some applications, may be prohibitive (**Figure 7A**). We evaluated Tronko using two different alignment methods, Needleman-Wunsch and Wavefront Alignment. In many cases, the two alignment algorithms perform similarly. However, in the case, where short, single-end reads are used (i.e., 150bp single-end reads), the Wavefront Alignment performs worse than the Needleman-Wunsch Alignment (see Figures S2D-F and S5D-F). The Wave-front Alignment algorithm implements heuristic modes to accelerate the alignment, which performs similar to Needleman-Wunsch when the two sequences being aligned are similar in length. However, when there is a large difference between the two sequences being aligned, we notice that the Wavefront Alignment forces an end-to-end alignment which contains large gaps at the beginning and end of the alignment. Hence, based on current implementations, we cannot recommend the use of the Wavefront Alignment for assignment purposes of short reads, although this conclusion could change with future improvements of the implementation of the wavefront alignment algorithm.

Tronko is currently not applicable to eukaryotic genomic data as it requires well-curated alignments of markers and associated phylogenetic trees, although we note that whole-genome phylogenetic reference databases for such data could potentially be constructed. Such extensions of the use of Tronko would require heuristics for addressing the memory requirements. Tronko currently has larger memory requirements than methods that are not phylogeny-based. Nonetheless, for assignment to viruses, amplicon sequencing and other forms of non-genomic barcoding, Tronko provides a substantial improvement over existing assignment methods and is the first full phylogenetic assignment method applicable to modern large data sets generated using Next Generation Sequencing.

The methods presented in this paper are implemented in the Tronko software package that includes Tronko-build and Tronko-assign for reference database building and species assignment, respectfully. Tronko can be downloaded at http://www.github.com/lpipes/tronko and is available under an open-software license.

## Methods

### Tronko-build reference database construction with a single tree

The algorithm used for assignment takes advantage of pre-calculated posterior probabilities of nucleotides at internal nodes of a phylogeny. We first estimate the topology and branch-lengths of the tree using RAxML^23^, although users of the method could use any tree estimation algorithm. We then calculate and store the posterior probabilities of each nucleotide in each node of the tree. For computational efficiency, this is done under a Jukes and Cantor (1969) model^24^, but the method can easily be extended to other models of molecular evolution. The calculations are achieved using an algorithm that traverses the tree only twice to calculate posterior probabilities simultaneously for all nodes in the tree. In brief, fractional likelihoods are first calculated in each node using a standard postorder traversal (e.g. Felsenstein 1981^25^). This directly provides the posterior probabilities in the root after appropriate standardization. An preorder traversal of the tree is then used to pull fractional likelihoods from the root down the tree to calculate posterior probabilities. While naive application of standard algorithms for calculating posterior probabilities in a node, to all nodes of a tree, have computational complexity that is quadratic in the number of nodes, the algorithm used here is linear in the number of nodes, as it calculates posterior probabilities for all nodes using a single postorder and a single preorder traversal without having to repeat the calculation for each node in the tree. For a single site, let the fractional likelihood of nucleotide *a ∈ {A, C, T, G}* in node *j* be *f_j_*(*a*), i.e., *f_j_*(*a*) is the probability the observed data in the site for all descendants of node *j* given nucleotide *a* in node *j*. Let *h_j_*(*a*) be the probability of the data in the subtree containing all leaf nodes that are not descendants of node *j*, given nucleotide *a* in node *j*, then the posterior probability of nucleotide *a* is (^26^):

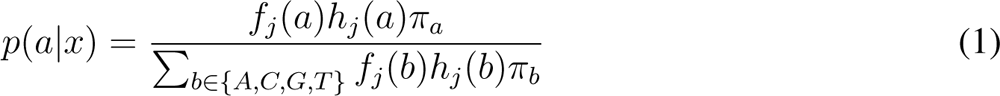

where *π_a_* is the stationary probability of nucleotide *a*. The algorithm here proceeds by first calculating and storing *f_j_*(*a*) for all values of *j* and *a* using a postorder traversal. It then recursively calculates *h_j_*(*a*) assuming time-reversibility using a preorder traversal as

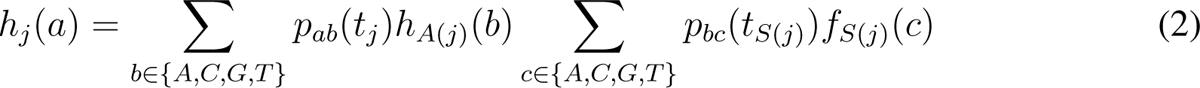

where *t_j_* is the branch length of the edge from node *j* to its parent, *p_ab_*(*t*) is the time dependent transition probability of a transition from nucleotide *a* to nucleotide *b* in time *t*, *A*(*j*) is the parent node of node *j* and *S*(*j*) is the the sister node of node *j* in the binary tree. The algorithm starts at the root with *h_root_*(*a*) = 1 *∀ a ∈ {A, C, T, G}* This algorithm is implemented in the program ‘Tronko-build’.

Each node in the tree is subsequently provided a taxonomy assignment. This is done by first making taxonomic assignments of the leaf nodes using the taxonomy provided by the taxid of the associated NCBI accession. We then make taxonomic assignments for internal nodes, at all taxonomic levels (species, genus, etc.), using a postorder traversal of the tree that assigns a taxonomic descriptor to node *i* if both children of node *i* have the same taxonomic assignment. Otherwise, node *i* does not have a taxonomic assignment at this taxonomic level and node *i* is given the next closest upwards taxonomic level where its children have the same taxonomic assignment. In other words, node *i* only gets a taxonomic assignment if the taxonomic assignments of both child nodes agree.

### Tronko-build reference database construction with multiple trees

MSAs for a large number of sequences can become unreliable, and computationally challenging to work with, due to the large number of insertions and deletions. For that reason, we devise an algorithm for partitioning of sequence sets into smaller subsets based on the accuracy of the alignment and using the inferred phylogenetic tree to guide the partitioning (**Figure S1**).

To measure the integrity of the MSA we calculate an average quality score, sum-of-pairs, *ASP* , which is a sum of pairwise alignment scores in the MSA. Assume a multiple sequence alignment of length *l* with *K* sequences, *A* = *{a_i,j_}*, where *a_i,j_* is the *j*th nucleotide in sequence *i*, 1 *≤ i ≤ K*, 1 *≤ j ≤ l*, *a_i,j_ ∈ M* = *{−, A, C, T, G, N }*. Define the penalty function, *p*:

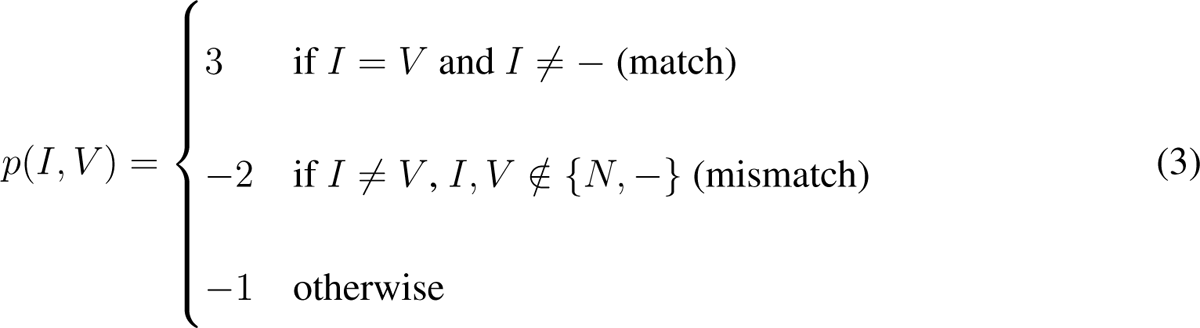

where *I, V ∈ M* . *ASP* is then calculated as

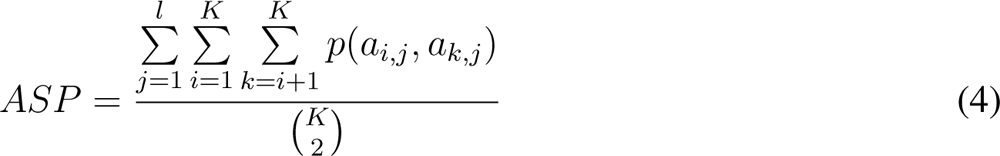

If the *ASP* is lower than the *ASP* threshold (a threshold of 0.1 was used in our analyses in this manuscript), the corresponding tree is split in three partitions at the node with the minimum variance, calculated as:

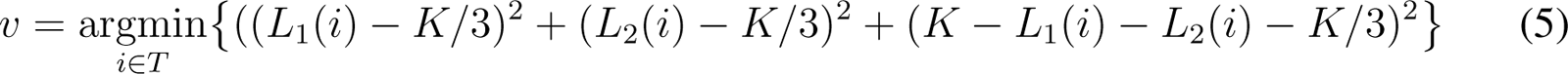

where *T* is a tree, i.e. a set of nodes, *L*_1_(*i*) and *L*_2_(*i*) is the number of leaf nodes descending from the left and right child node, respectively, of node *i*, and *K* is the total number of leaf nodes in the tree. We then split the tree into 3 subtrees by eliminating node *v*. Each partition is realigned with `FAMSA`^27^ and new trees are constructed using `RAxML`^23^ using default parameters and the GTR+Gamma model.`FAMSA` is used to optimize for speed since it is 1 to 2 orders of magnitude faster than Clustal^28^ or MAFFT^29^ with similar quality (see Deorowicz et al., 2016). We explored different combinations of tree estimation methods (including IQ-TREE2^30^), multiple sequence aligners, and global aligners (**Figure S3**). While most combinations of methods were quite similar (especially for the genus level), the use of FAMSA+RAxML+NW was optimal with regards to speed and accuracy. We ran IQ-TREE2 with the default settings using options `-m GTR+G -nt 4` to be consistent with similar RAxML settings. The sequences are recursively partitioned until the *ASP* score is above the threshold. Finally, the trees, multiple sequence alignments, taxonomic information, and posterior probabilities are written to one reference file which can be loaded for subsequent assignment of reads. Notice, that the procedure for phylogeny estimating and calculation of posterior probabilities only has to be done once for a marker and then can be used repeatedly for assignment using different data sets of query sequences.

### Simulation of query sequences

To simulate single-end reads from a reference sequence, a starting point is selected uniformly at random and extends for *m*_0_ base pairs, where *m*_0_ represents the read length. For paired-end reads a similar random selection of a starting point occurs, extending *m*_0_ base pairs. From the end of this read, if the insert size *m*_1_ is positive, the reverse read begins *m*_1_ base pairs forward with a length of *m*_0_. If *m*_1_ is negative, the reverse read starts *m*_1_ base pairs backward. Sequencing errors are then added independently with different probabilities *α* = 0, *α* = 0.01, and *α* = 0.02 at each site. These errors are induced by changing the nucleotide to any of the other three possible nucleotides, following the probabilities used by Stephens et al. (2016) ^31^:

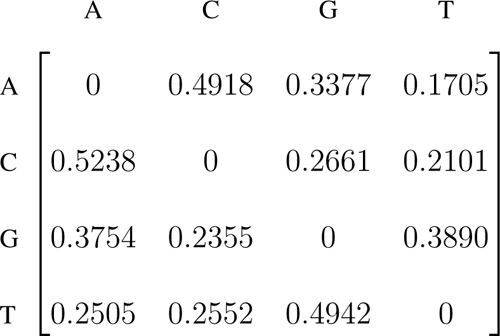

### Taxonomic classification of query sequences

First, `BWA-MEM`^32^ is used with default options to align the query sequences to the reference sequences, thereby identifying a list of the highest scoring reference sequences (which we designate as BWA-MEM hits) from the reference database. We use `BWA-MEM` as the original Minimap2 manuscript^33^ demonstrated that `BWA-MEM` had the lowest error rate for the same amount of fractional mapped reads when compared to Minimap2, SNAP^34^, and bowtie2^35^. Second, a global alignment, either using the Needleman-Wunsch algorithm^14^ or the Wavefront alignment algorithm^15^, is performed only on the sequence with the highest score from each subtree (reference sequence set) identified using the previously described partitioning algorithm.

Once aligned to the reference sequence, a score, *S*(*i*) is calculated for all nodes, *i*, in the tree(s) that the reference sequence is located to. For a given read, let *b_j_* be the observed nucleotide in the position of the read mapping to position *j* in the alignment. We also assume an error rate, *c*. For example, if the true base is G and the error rate is *c*, then the probability of observing A in the read is *c/*3. We note that this error rate can be consider to include both true sequencing errors and polymorphisms/sequence divergence. In an ungapped alignment, the score for site *j* in node *i* is then the negative log of a function that depends on the posterior probability of the observed nucleotide in the query sequence, IP*_ij_*(*b_j_*), and the error rate:

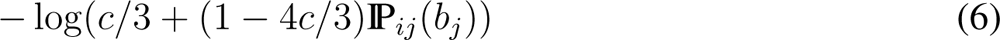

Assuming symmetric error rates, the probability of observing the base by error is (1 *−* IP*_ij_*(*b_j_*))*c/*3 and the probability of observing the base with no error is (1 *− c*)IP*_ij_*(*b_j_*). The sum of these two expressions equals the expression in the logarithm above. The score for all *s* sites in the read is defined as 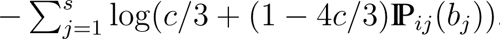.

Notice that the full phylogenetic likelihood for the entire tree, under standard models of molecular evolution^26^ with equal base frequencies and not accounting for errors, and assuming time reversibility, is

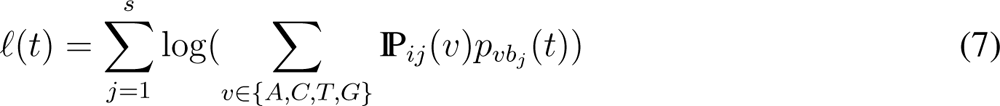

where *p_vb__j_* (*t*) is the time dependent transition probability from base *v* to base *b_j_* in time *t*. This statement takes advantage of the fact that, under time-reversibility, the posterior for a base in an node is proportional to the fractional likelihood of that base in the node, if the tree is rooted in the node. For small values of *t*, *t* converges to log(IP*_ij_*(*b_j_*). Minimizing the score function, therefore, corresponds to maximizing the full phylogenetic likelihood function assuming that the branch leading to the query sequence is infinitesimally short and connects with the tree in an existing node. An alternative interpretation is that the score maximizes the probability of observing the query sequence if it is placed exactly in a node or, equivalently, minimizes the expected mismatch between the query and a predicted sequence sampled form the node.

To address insertions and deletions, we define scores of *γ* and *λ* for a gap or insertion, respectively, in the query sequence relative to the reference sequence. We also entertain the possibility of a gap in the reference sequence in node *i* in read position *j*, *r_ij_*, which occurs when the reference is a leaf node with a gap in the position or if it is an internal node with all descendent nodes having gaps in the position. We use the notation *M_g_* = *{−, N }* for gaps and *M_n_* = *{A, C, T, G}* for nucleotides (no gap). Then, the score for node *i* in site *j* of the read, with observed base *b_j_*, is

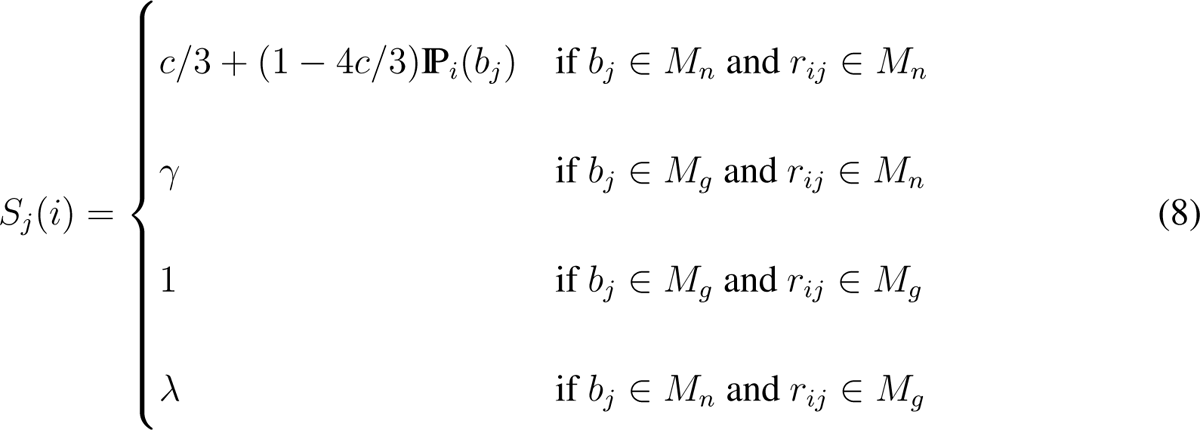

The total score for the entire read is

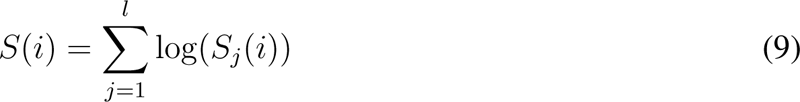

For paired reads, the scores for each node in the tree is calculated as the sum of the scores for the forward read and the scores for the reverse read. Scores are calculated for all nodes in each tree that contain a best hits from the `bwa mem` alignment. For all analyses in this paper we use values of *c* = 0.01, *λ* = 0.01, and *γ* = 0.25.

After calculation of scores, the LCA of all of the lowest scoring nodes, using a user-defined cut-off parameter, is calculated. For example, if the cut-off parameter is 0, only the highest scoring node (or nodes with the same score as the highest scoring node) is used to calculate the LCA. If the cut-off parameter is 5, the highest scoring node along with all other nodes within a score of 5 of the highest scoring node are used to calculate the LCA. Once the LCA node is identified, the classification of the single read (or paired-reads) will be assigned to the taxonomy assigned to that node. The classification of query sequences is parallelized.

### Taxonomic assignment using pplacer

To generate phylogenetic placements using pplacer, we first aligned sequencing reads to the reference sequences using hmmer3^36^. We then ran pplacer, rppr prep db, and guppy classify all using the default parameters in that order. Next, to obtain taxonomic assignments, we used the R package BoSSA^37^ to merge the multiclass element (which is a a data frame with the taxonomic assignments of each placement) and the placement table of pplace object (the output of pplacer) and only kept the “best” type of placement for each read. For paired-end sequences, we assigned the taxonomy by the LCA of both pairs of reads.

### Taxonomic assignment using APPLES-2

To generate phylogenetic placements using APPLES-2. We first aligned sequencing reads to the reference sequences using hmmer3^36^. We then converted the alignment output from Stockholm to FASTA format and then separated the reference sequences from the sequencing reads (an input requirement for APPLES-2) using in-house scripts. We then ran `run apples.py` with the default parameters. In order to ensure that the tree that was output from APPLES-2 was strictly binary (a requirement to assign taxonomy), we extracted the tree from the jplace output and resolved polytomies using the `multi2di` function from the R package ape^38^. Next, we ran `run apples.py` again using the output tree from ape (with option `--tree=`) and disabled reestimation of branch lengths (in order to keep the tree as strictly binary) by using the option `--disable-reestimation`. To assign taxonomy we ran `gappa examine assign` from the Gappa toolkit^39^ using the options `--per-query-results` and `--best-hit`.

### Classification metrics used for accuracy evaluations

We used the taxonomic identification metrics from Siegwald *et al.* 2017^40^ and Sczyrba *et al.*2017^41^. A true positive (TP) read at a certain taxonomic rank has the same taxonomy as the sequence it was simulated from. A misclassification (FP) read at a certain taxonomic rank has a taxonomy different from the sequence it was simulated from. A false negative (FN) read, at a certain taxonomic rank, is defined as a read that received no assignment at that rank. For accuracy, we use the following measures for recall and misclassification rate.

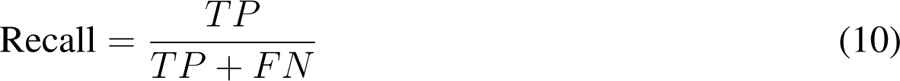

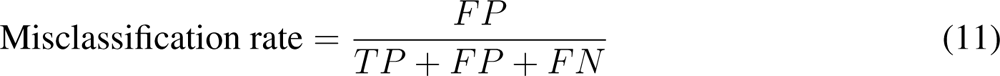

### Classification of mock community reads

For Schirmer *et al.* (2015) we used the ERR777705 sample, for Gohl *et al.* (2016) we used the SRR3163887 sample, and for Braukmann *et al.* (2019) we used the SRR8082172 sample. For Lluch *et al.* (2015)^22^ we used the ERR1049842 sample. All sample raw reads used for assignment were first filtered through the Anacapa Quality Control pipeline^16^ with default parameters up until before the amplicon sequence variant (ASV) construction step. Only paired reads were retained for assignment. For mock datasets where the true species were only defined with “sp.”, species assignment were excluded for all methods. After Tronko assignments, we filtered results using a script to check the number of mismatches in the forward vs. reverse reads, and used a *χ*^2^-distribution to filter out assignments that have a discrepancy in mismatches.

### Leave-species-out and leave-one-individual-out analyses

We used two datasets (Charadriiformes and Bacteria) for leave-species-out and leave-one-individual-out analyses. For one dataset, we used 1,467 COI reference sequences from 253 species from the order Charadriiformes. For the leave-species-out analyses with Charadriiformes we removed each of the species one at a time (excluding singletons, i.e. species only represented by a single sequence), yielding 252 different reference databases. For the leave-species-out analyses with Bacteria, we randomly selected 5, 000 taxonomically divergent bacteria species from the 16S reference database built through CRUX. For the leave-species-out analyses with Bacteria, we removed each of the species, one at a time (excluding singletons), yielding 2, 323 different reference databases. For each database, we then simulated reads from the species that had been removed with different error rates, and assigned to taxonomy using all methods tested (Tronko, kraken2, metaphlan2, MEGAN, pplacer, and APPLES-2), using the same reference databases and same simulated reads for all methods. For the leave-individual-out analysis with Charadriiformes, we removed a single individual from each species (excluding singletons) yielding 1,423 different reference databases. Assignments for all method were performed with default parameters and where a paired read mode was applicable, that mode was used when analyzing paired reads. For paired-end read assignments with MEGAN, the assignment is the LCA of the forward and reverse read assignments as described in the MEGAN manual v6.12.3. For metaphlan, the results from the forward reads and reverse reads were combined.

### Custom 16S and COI Tronko-build reference database construction

For the construction of the reference databases in this manuscript, we use custom built reference sequences that were generated using common primers^42–45^ for 16S and COI amplicons that have been used in previous studies^46–48^ using the CRUX module of the Anacapa Toolkit^16^. For the COI reference database, we use the following forward primer: GGWACWGGWTGAACWGTWTAY-CCYCC, and reverse primer: TANACYTCnGGRTGNCCRAARAAYCA from Leray *et al.*(2013) and Geller *et al.* (2013)^43, 44^, respectively, as input into the CRUX pipeline^16^ to obtain a fasta and taxonomy file of reference sequences. For the 16S database, we use forward primer: GT-GCCAGCMGCCGCGGTAA, and reverse primer: GACTACHVGGGTATCTAATCC from Capo-raso *et al.* (2012)^42^. We set the length of the minimum amplicon expected to 0bp, the length of the maximum amplicon expected to 2000bp, and the maximum number of primer mismatches to 3 (parameters `-s 0`, `-m 2000`, `-e 3`, respectively). Since all of the custom built libraries contain *≥*500,000 reference sequences and MSAs, we first used Ancestralclust ^49^ to do an initial partition of the data, using parameters of 1000 seed sequences in 30 inital clusters (parameters `-r 1000` and `-b 30`, respectively). For the COI database, we obtain 76 clusters and for the 16S database we obtain 228 clusters. For each cluster, we use FAMSA^27^ with default parameters to construct the MSAs and RAxML^23^ with the model GTR+Γ of nucleotide substitution to obtain the starting trees for Tronko-build. The identified reference databases, MSAs, phylogeentic trees, and posterior probabilites of nucleotides in nodes for COI and 16S, are available for download at https://doi.org/10.5281/zenodo.13182507.

## Acknowledgements

We would like to thank Rachel Meyer and CALeDNA for their support in this project. We acknowledge Thorfinn Sand Korneliussen for advice on parallelization of the method.

## Inclusion and diversity

We support inclusive, diverse, and equitable conduct of research.

## Supplementary Material

**Figure S1:**
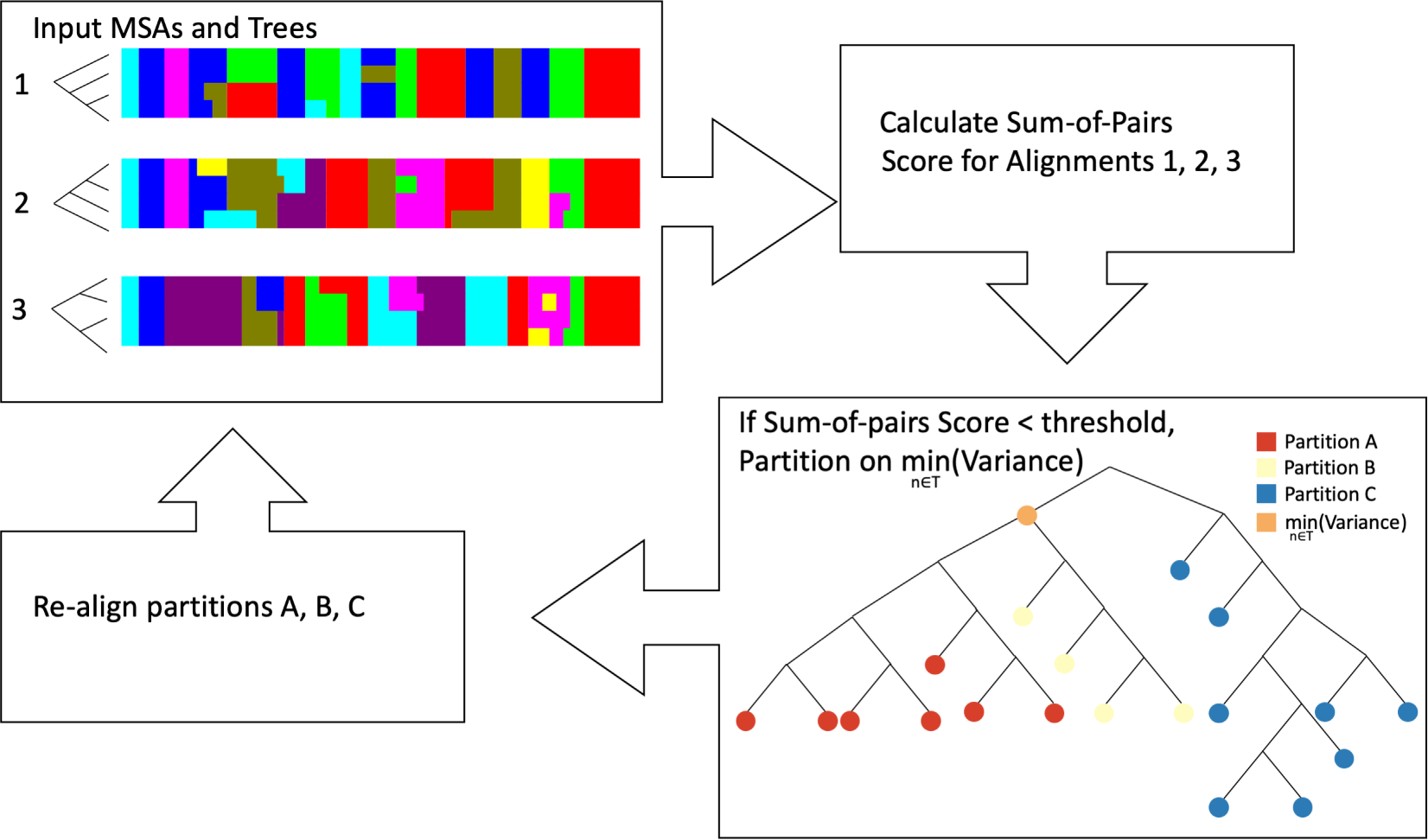
Workflow of iterative partitioning procedure. First, the MSAs and corresponding phylogenetic trees are used as input into the algorithm. Then, the sum-of-pairs scores are calculated for each partition. If the sum-of-pairs score is below a heuristic threshold, the tree is used to partition the sequences into three partitions in the cluster based on the node with the smallest variance. Each of the three partitions, is re-aligned and phylogenetic trees are estimated. The algorithm stops for a given partition when the sum-of-pairs score is greater than the heuristic threshold.

**Figure S2:**
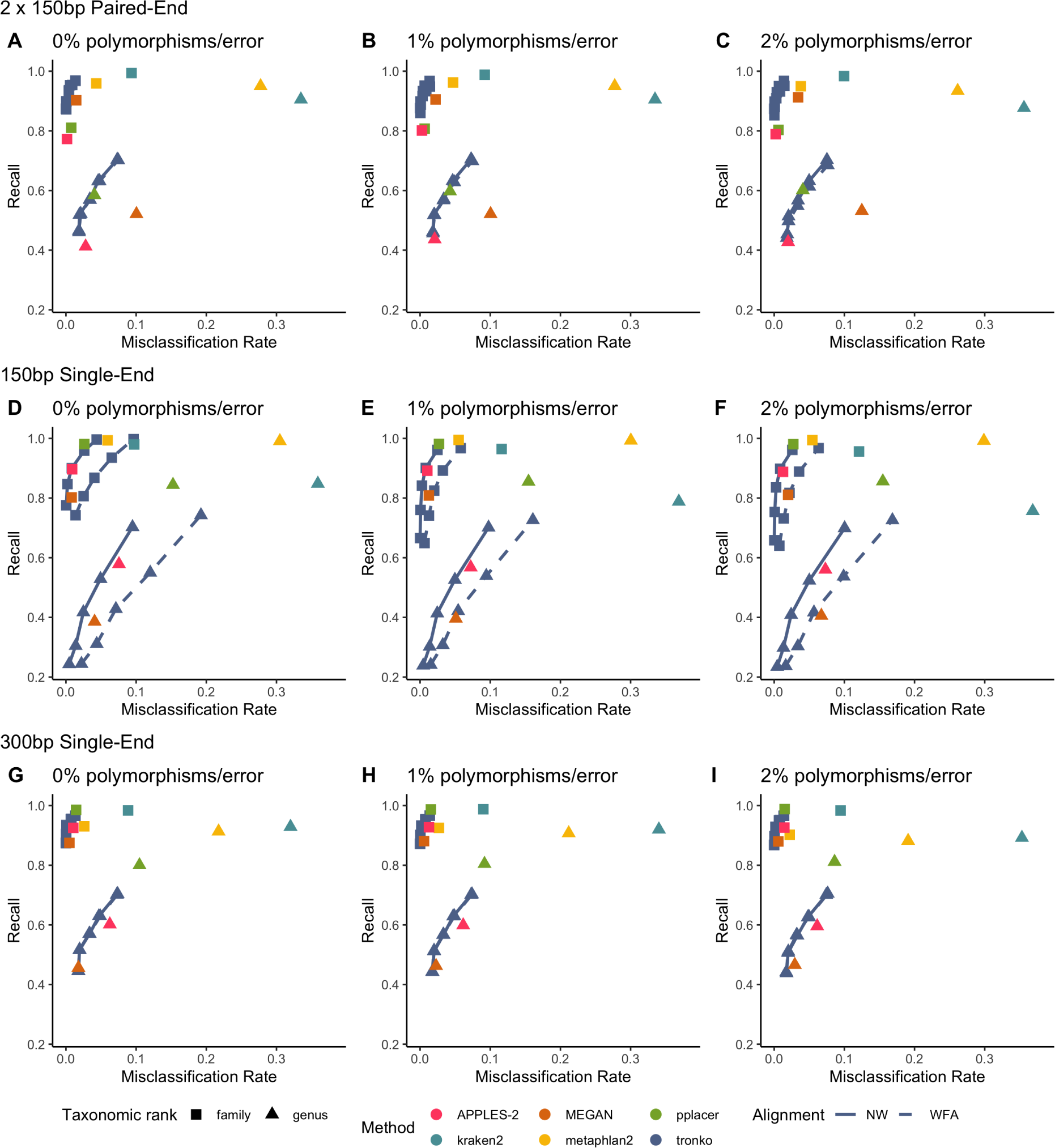
Recall vs. Misclassification rates using leave-one-species-out analysis for the order Charadriiformes (COI metabarcode) with paired-end 150bp*×*2 reads with 0% (A), 1% (B), and 2% (C) error/polymorphism, single-end 150bp reads with 0% (D), 1% (E), and 2% (F) error/polymorphism, and single-end 300bp reads with 0% (G), 1% (H), and 2% (I) error/polymorphism using kraken2, metaphlan2, MEGAN, pplacer, APPLES-2, and Tronko with cut-offs of 0, 5, 10, 15, and 20 using the Needleman-Wunsch alignment (solid line) and Wavefront alignment (dashed line).

**Figure S3:**
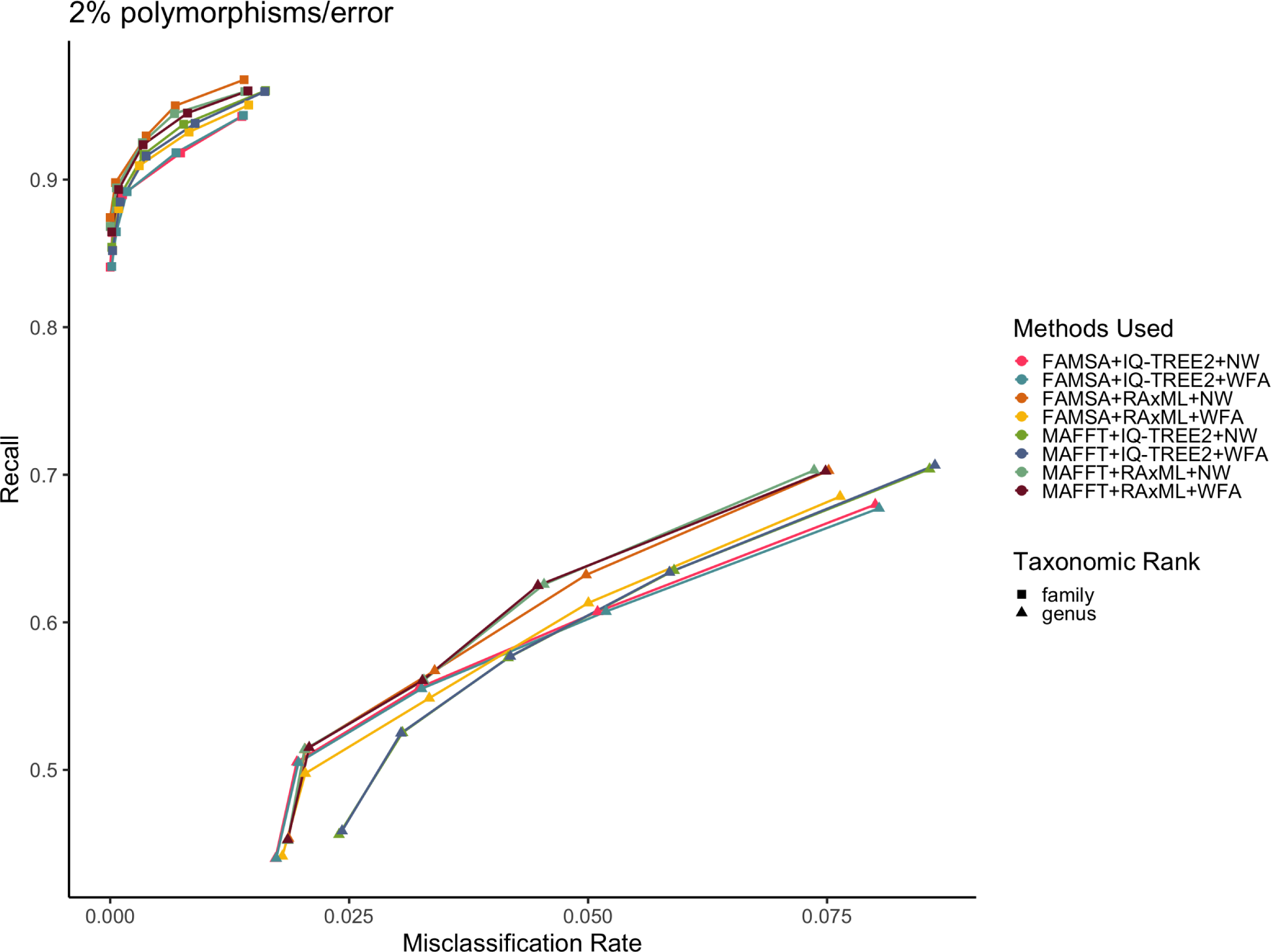
Recall vs. Misclassification rates using leave-one-species-out analysis for the order Charadriiformes (COI metabarcode) with paired-end 150bp*×*2 reads with 2% error/polymorphism using Tronko with cut-offs of 0, 5, 10, 15, and 20 and different combinations of tree estimation methods and aligners. For tree estimation we used RAxML and IQ-TREE2. For multiple sequence aligners, we used FAMSA and MAFFT. For global alignment methods, we used Needleman-Wunsch (NW) and Wavefront Alignment (WFA). Colors represent different combinations of methods.

**Figure S4:**
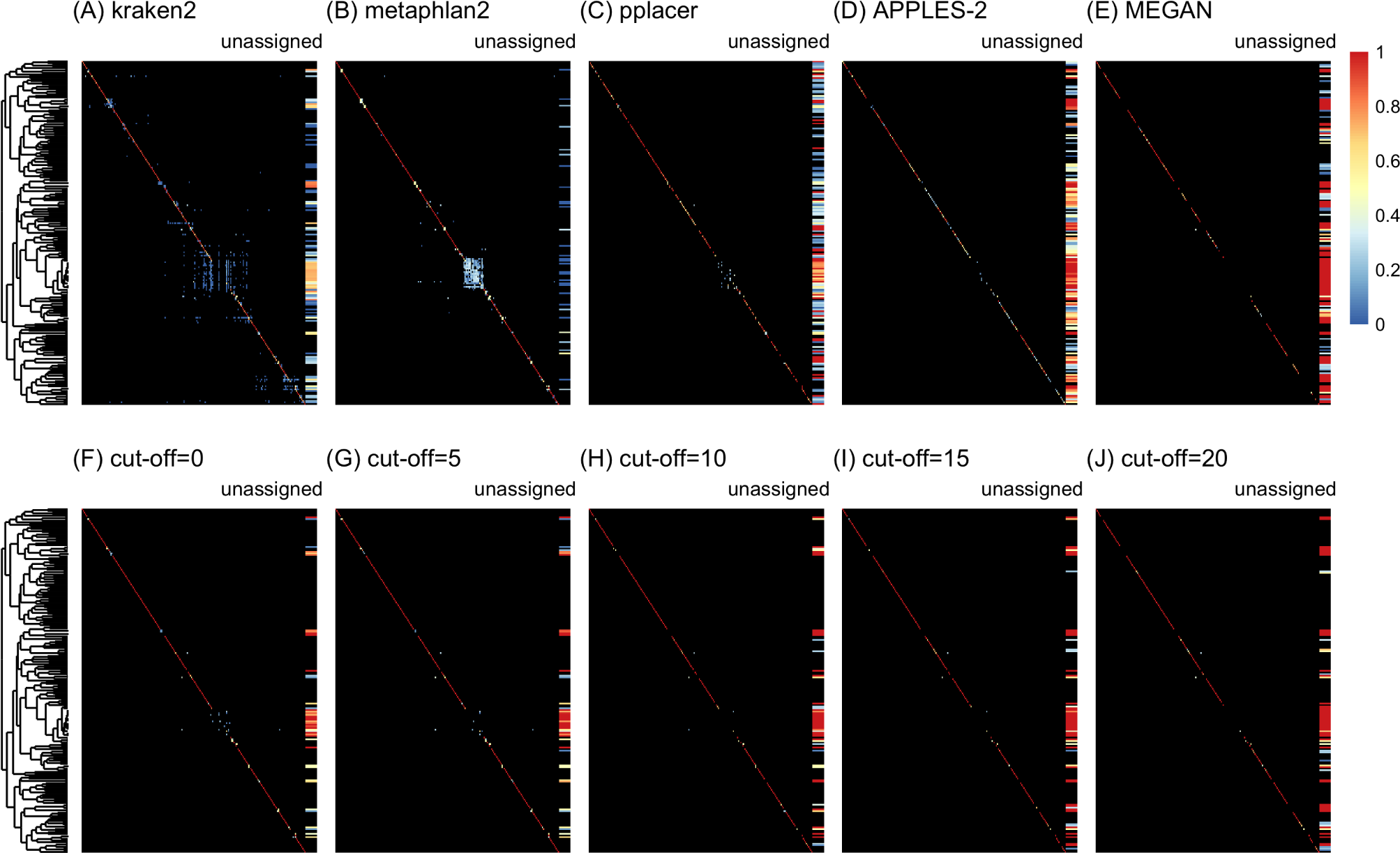
Confusion matrices at the species level of the order Charadriiformes using the leave-one-individual-out analysis with paired-end 150bp*×*2 reads with 2% error/polymorphism using kraken2 (A), metaphlan2 (B), pplacer (C), APPLES-2 (D), MEGAN (E), and Tronko using the Needleman-Wunsch alignment (NW) for cut-offs 0 (F), 5 (G), 10 (H), 15 (I), and (J) 20. Unassigned column contains both unassigned queries and queries assigned to a lower taxonomic level. Phylogenetic tree represents ancestral sequences at the species level.

**Figure S5:**
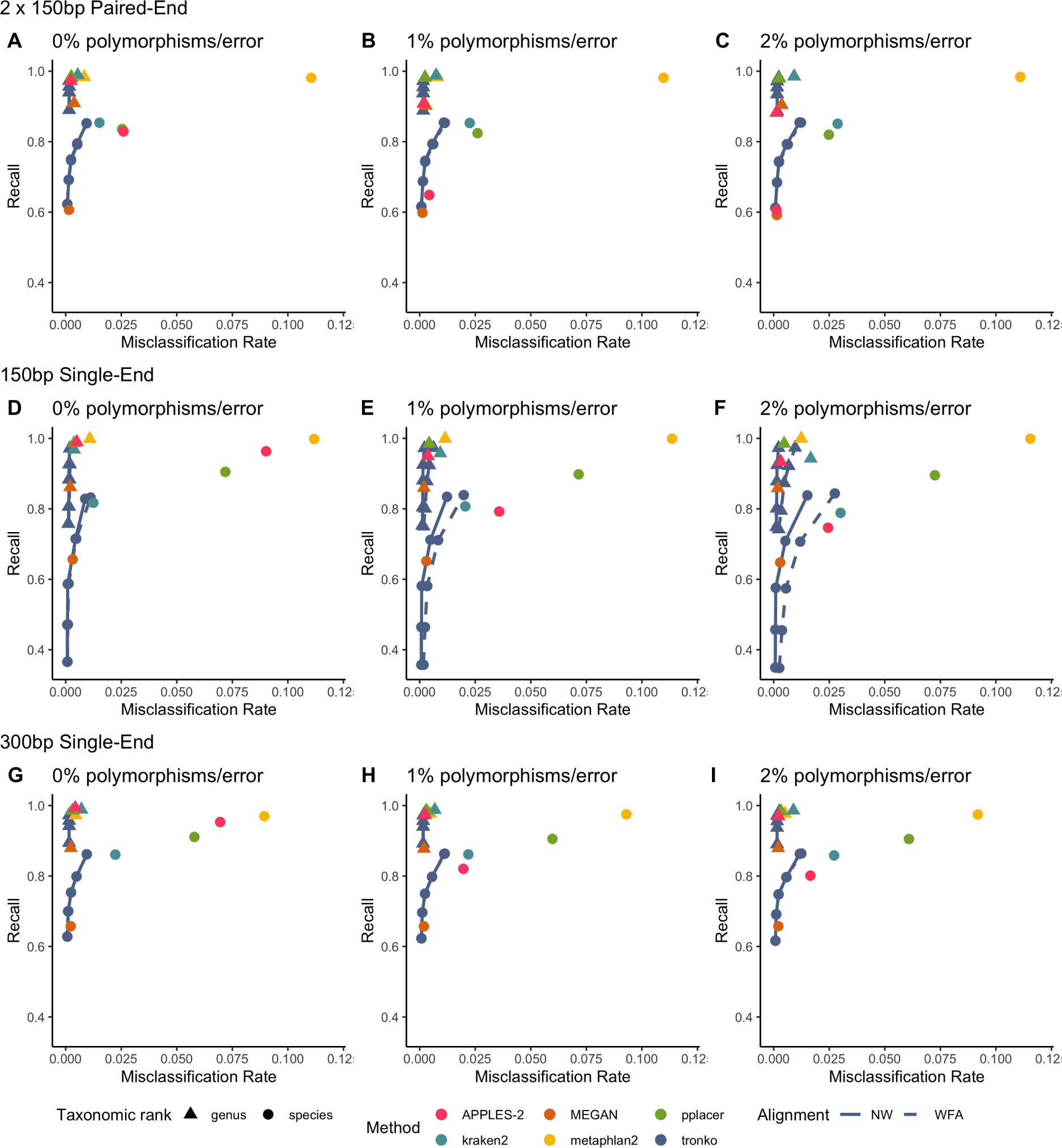
Recall vs. Misclassification rates using leave-one-individual-out analysis for the order Charadriiformes (COI metabarcode) with paired-end 2*×*150bp reads with 0% (A), 1% (B), and 2% (C) error/polymorphism, single-end 150bp reads with 0% (D), 1% (E), and 2% (F) error/polymorphism, and single-end 300bp reads with 0% (G), 1% (H), and 2% (I) error/polymorphism using kraken2, metaphlan2, MEGAN, pplacer, APPLES-2, and Tronko with cut-offs of 0, 5, 10, 15, and 20 using the Needleman-Wunsch alignment (solid line) and Wavefront alignment (dashed line).

**Figure S6:**
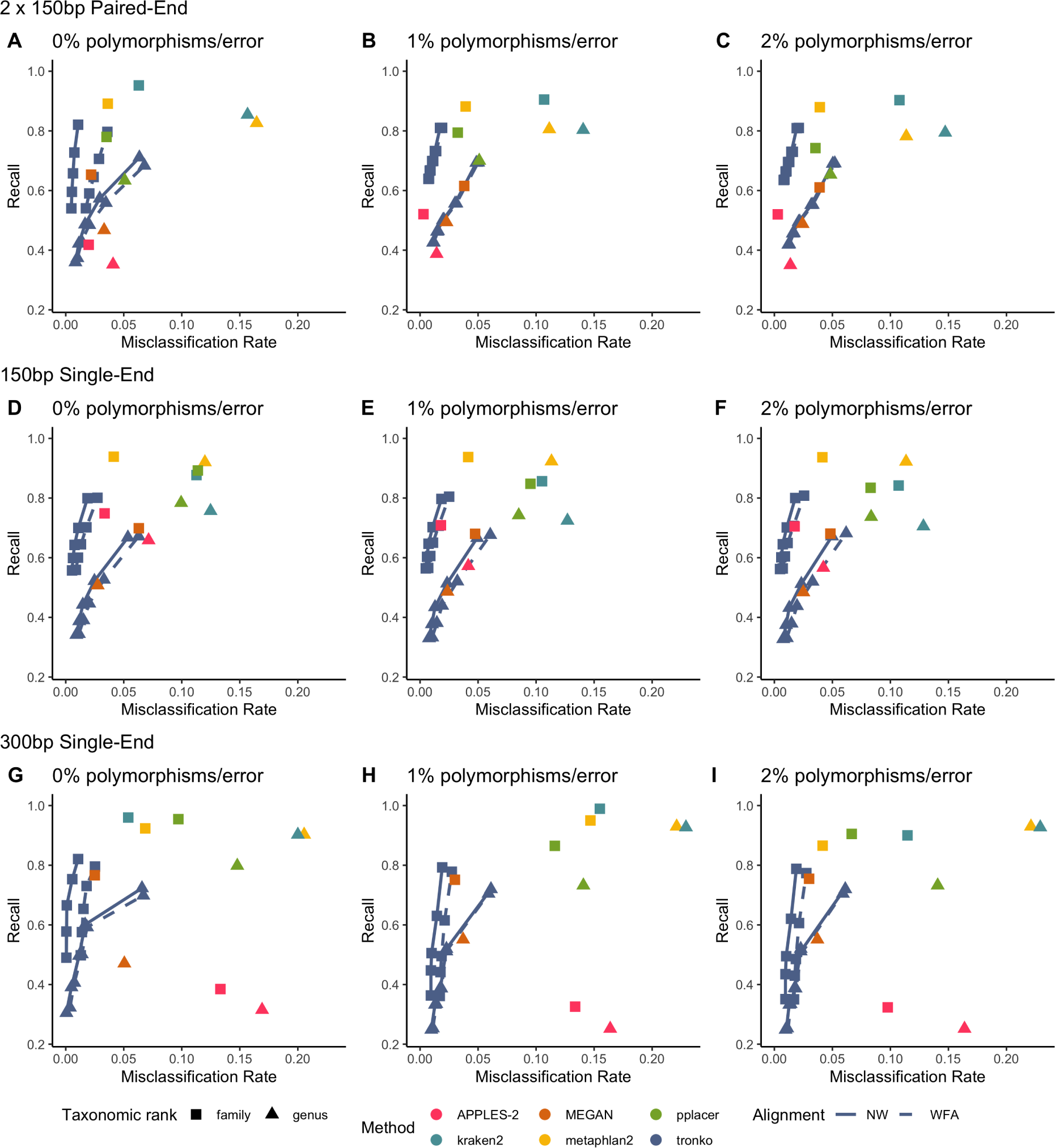
Recall vs. Misclassification rates using leave-one-individual-out analysis for the order Charadriiformes (COI metabarcode) with paired-end 2*×*150bp reads with 0% (A), 1% (B), and 2% (C) error/polymorphism, single-end 150bp reads with 0% (D), 1% (E), and 2% (F) error/polymorphism, and single-end 300bp reads with 0% (G), 1% (H), and 2% (I) error/polymorphism using kraken2, metaphlan2, MEGAN, pplacer, APPLES-2, and Tronko with cut-offs of 0, 5, 10, 15, and 20 using the Needleman-Wunsch alignment (solid line) and Wavefront alignment (dashed line).

**Figure S7:**
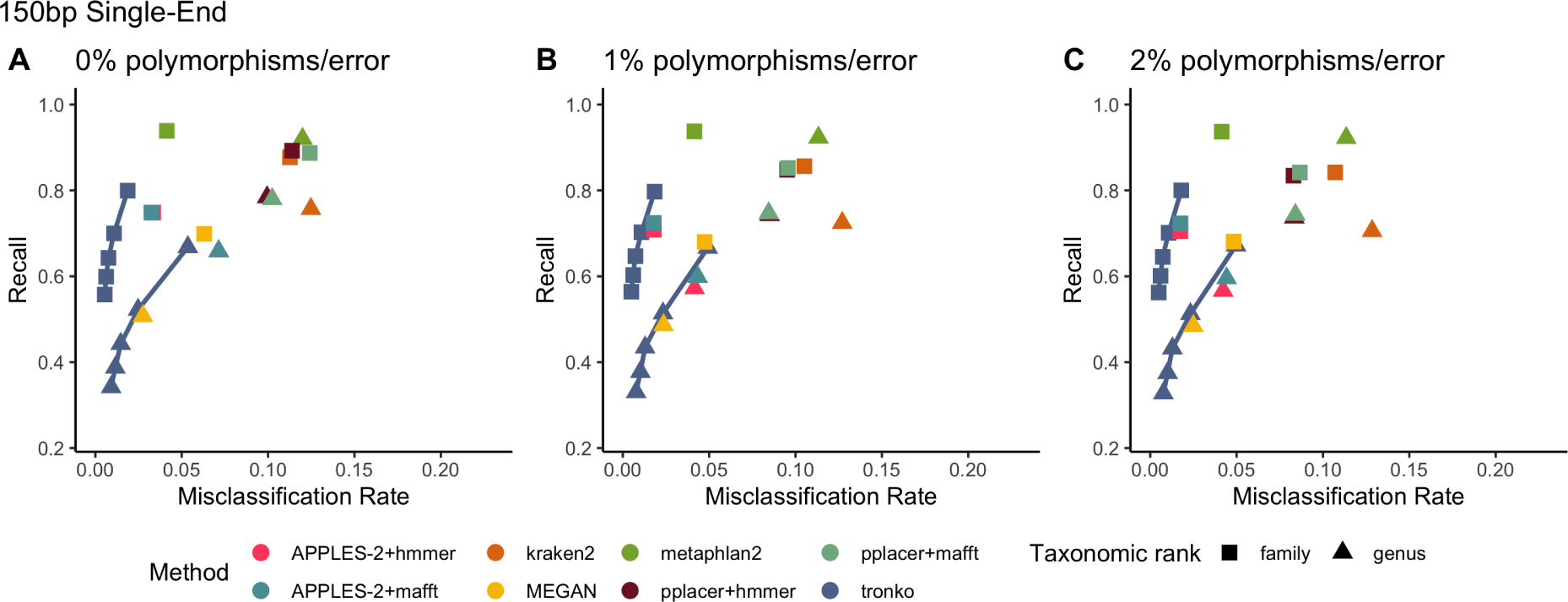
Recall vs. Misclassification rates using leave-one-individual-out analysis for bacterial species (16S metabarcode) with paired-end 2*×*150bp reads with 0% (A), 1% (B), and 2% (C) error/polymorphism using kraken2, metaphlan2, MEGAN, pplacer+hmmer, pplacer+mafft, APPLES-2+hmmer, APPLES-2+mafft, and Tronko with cut-offs of 0, 5, 10, 15, and 20 using the Needleman-Wunsch alignment (solid line).

**Figure S8:**
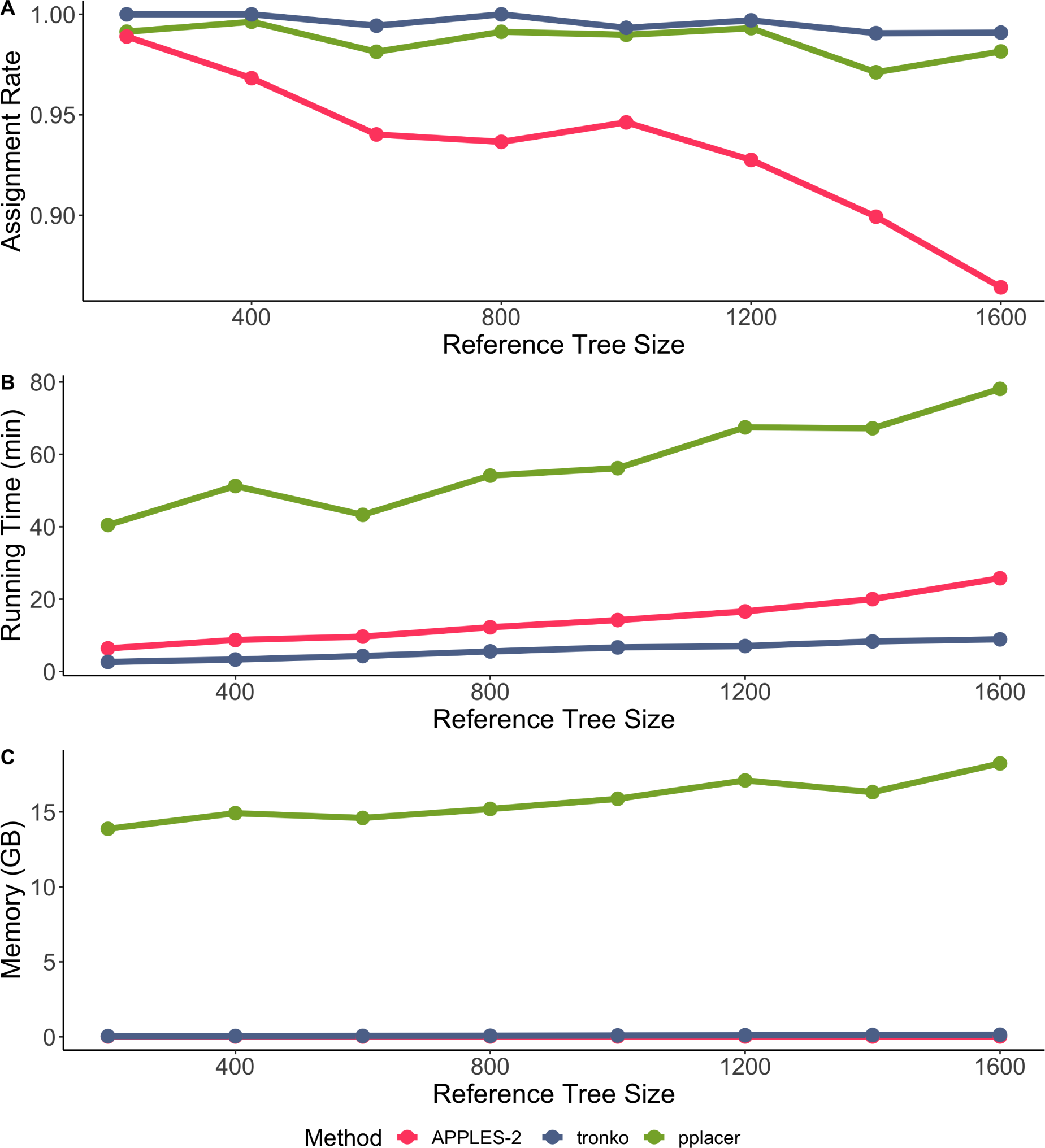
Comparisons of Tronko with pplacer and APPLES-2 using a database of 200, 400, 600, 800, 1000, 1200, 1400, and 1600 reference sequences. (A) Assignment rate against the number of reference sequences at the species level. (B) Running time against the number of reference sequences. (C) Peak memory in gigabytes against number of references. Both methods had a 100% true positive rate for all sizes of databases. Assignment rate is the number of reads assigned at the species level for each method. Reference sequences were chosen randomly from the COI reference database.

**Figure S9:**
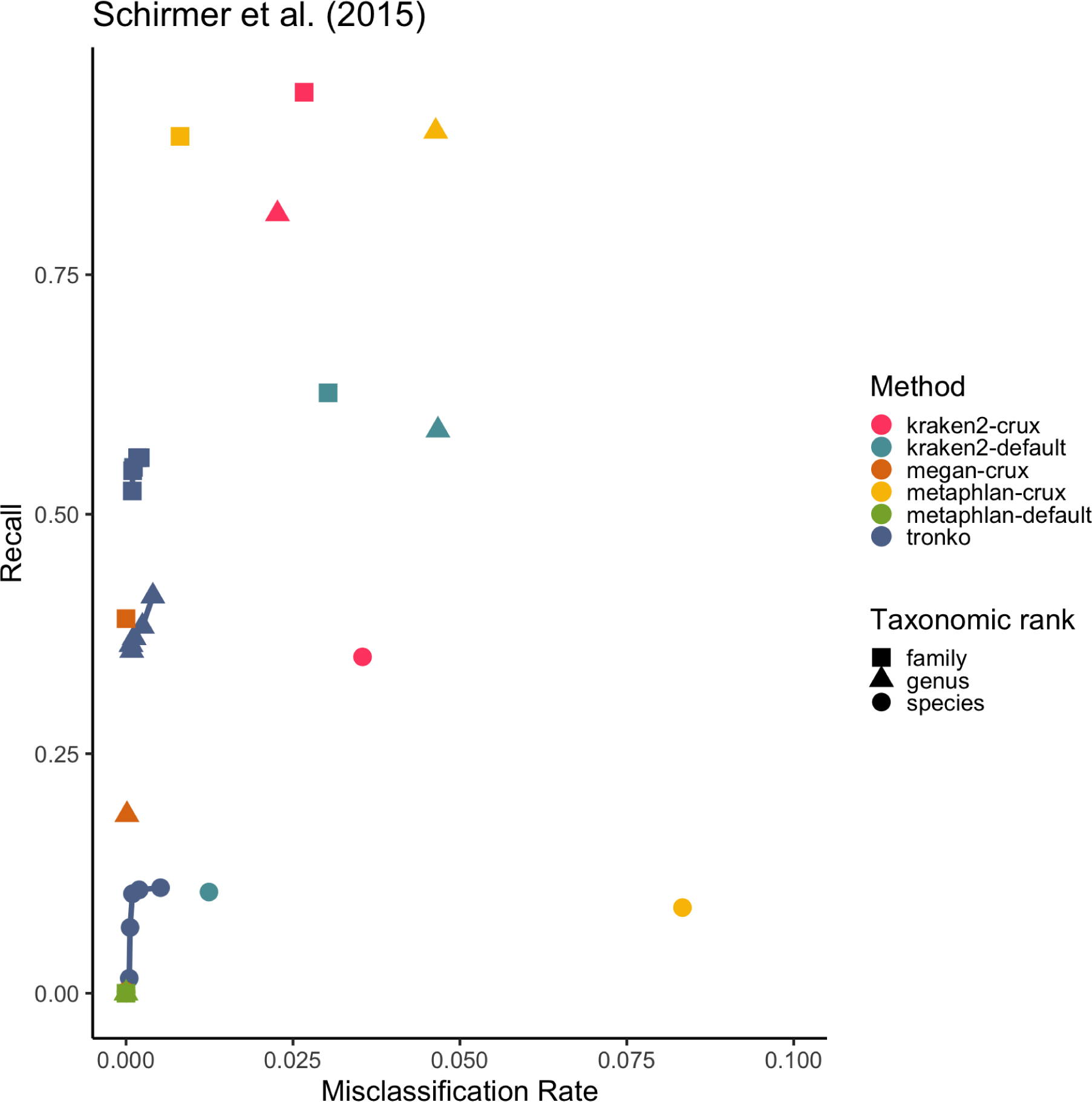
Close-up figure of Figure 6A.

**Figure S10:**
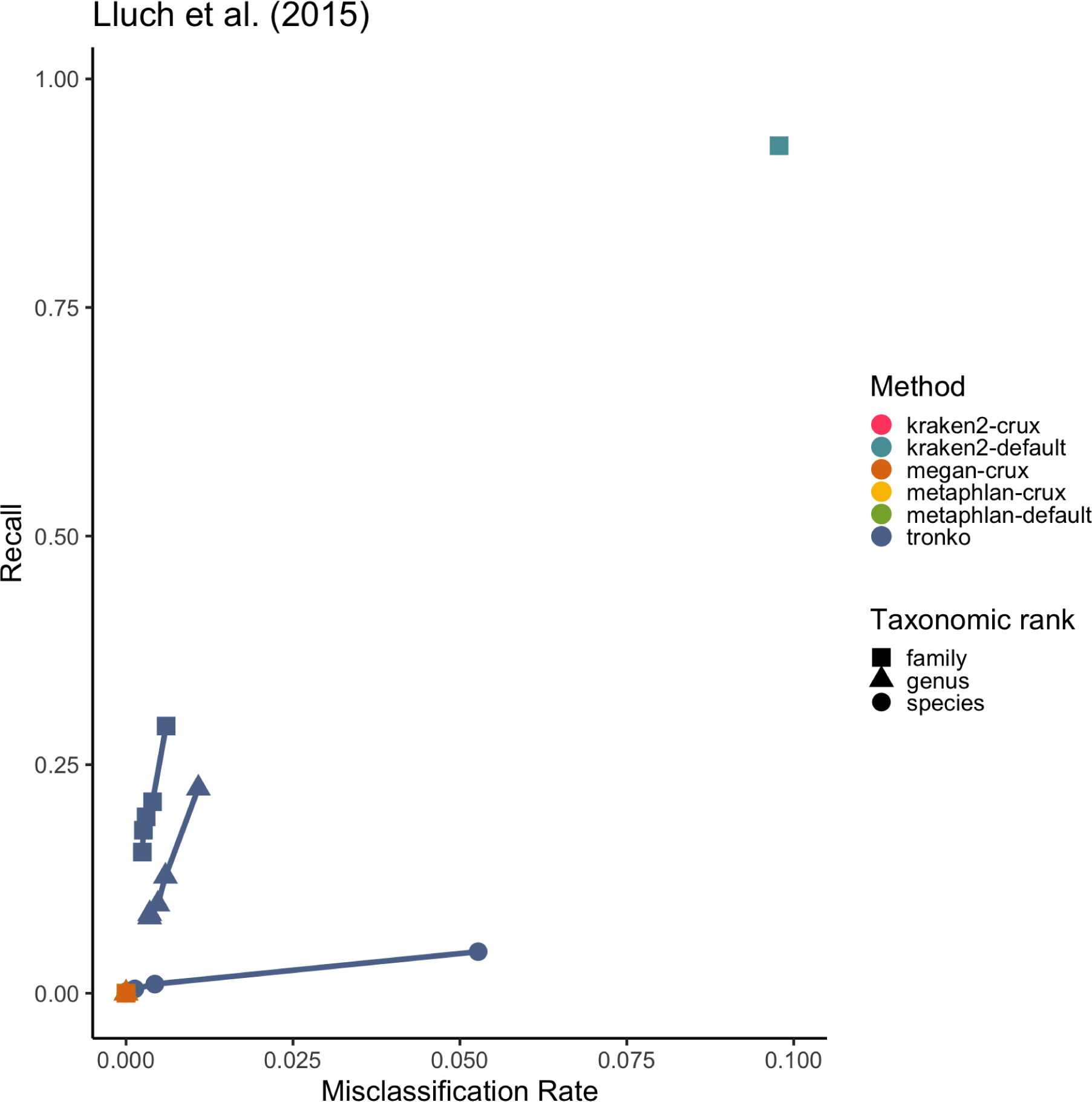
Close-up figure of Figure 6A.

**Figure S11:**
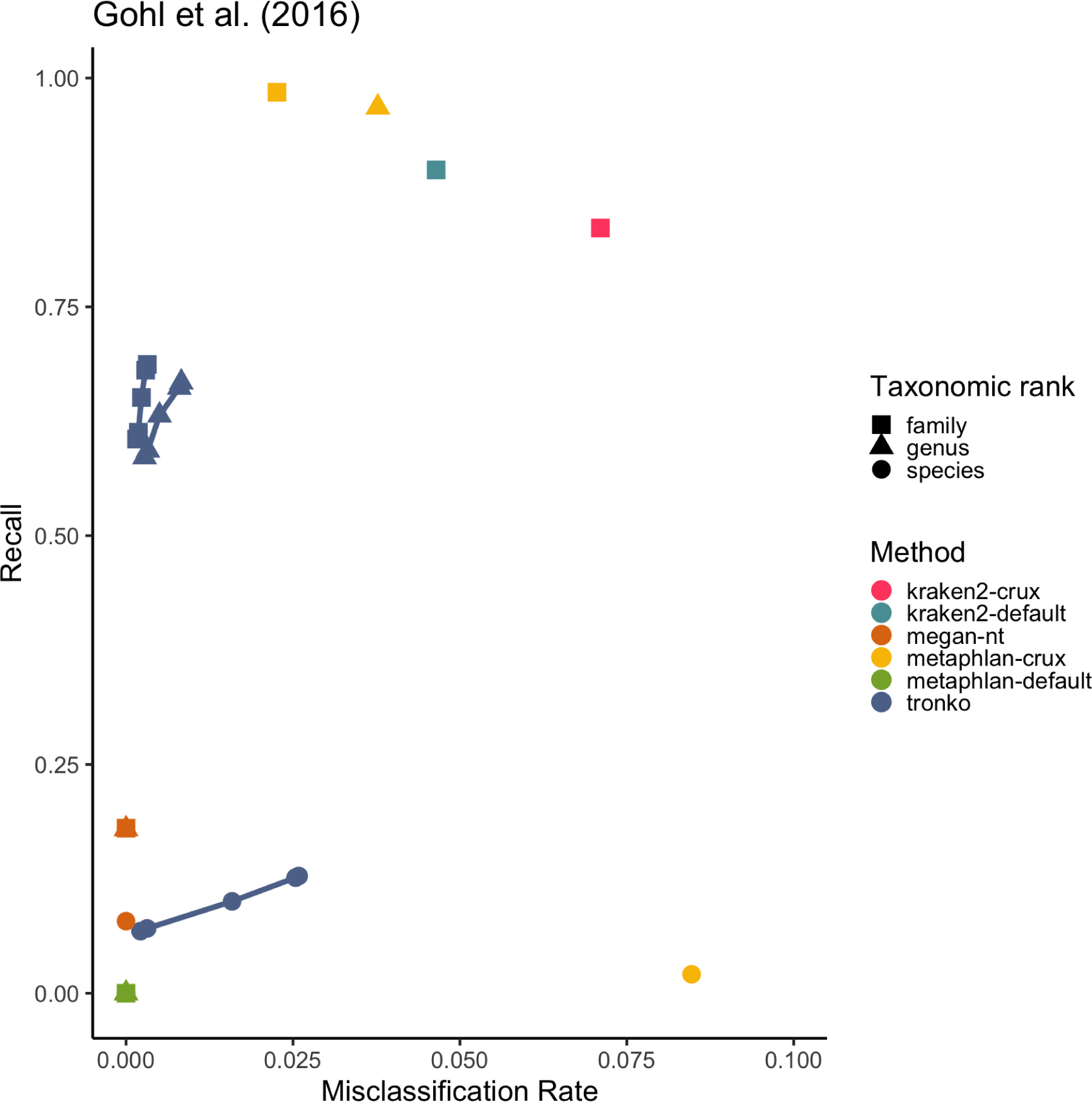
Close-up figure of Figure 6B.

**Figure S12:**
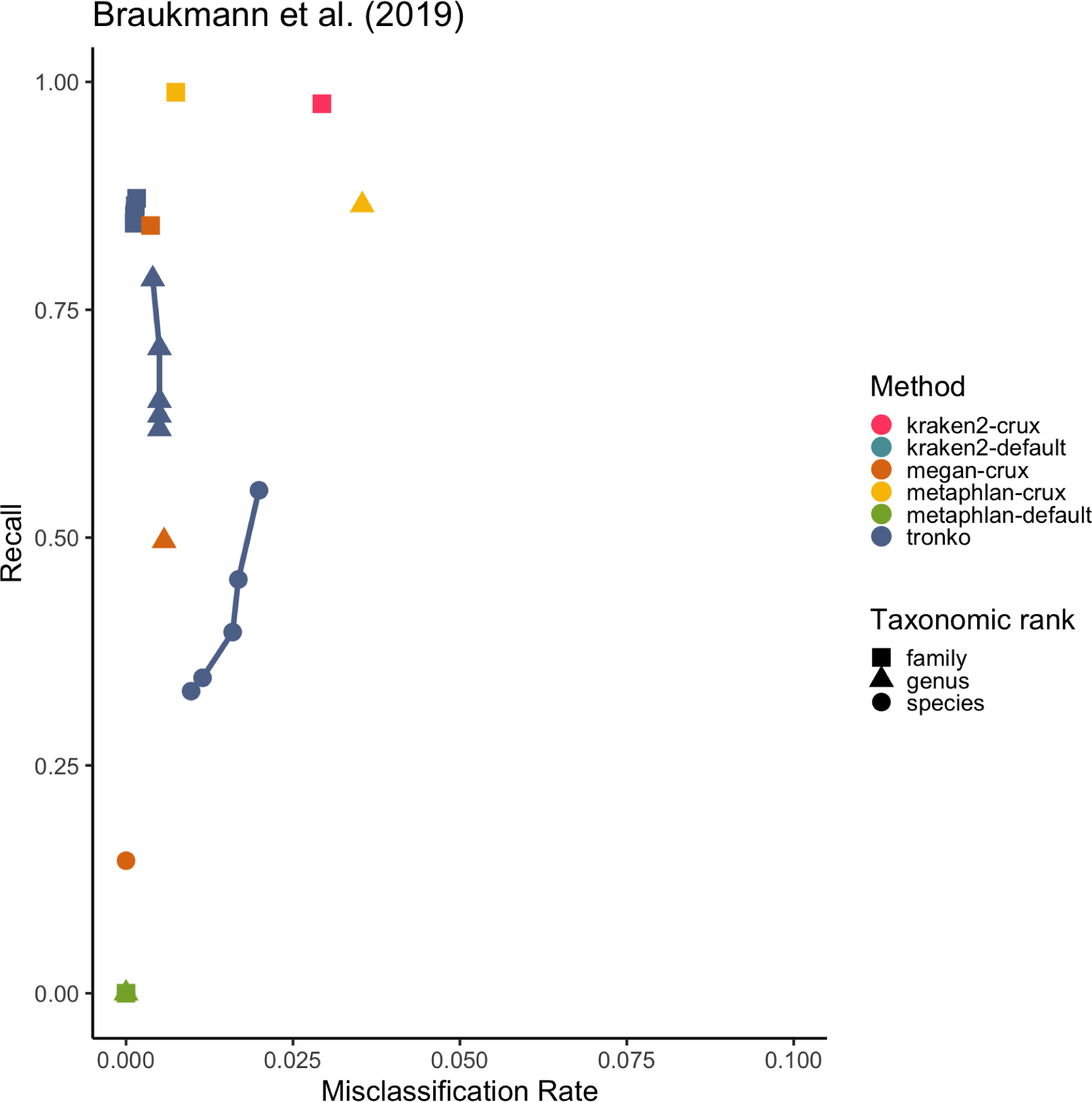
Close-up figure of Figure 6C.

